# Regulation of intracellular signaling and neuron function by Bardet-Biedl Syndrome proteins in patient-specific iPSC-derived neurons

**DOI:** 10.1101/2020.10.29.359851

**Authors:** Liheng Wang, Yang Liu, George Stratigopoulos, Sunil Panigrahi, Lina Sui, Charles A. Leduc, Hannah J. Glover, Maria Caterina De Rosa, Lisa C. Burnett, Damian J. Williams, Linshan Shang, Robin Goland, Stephen H. Tsang, Sharon Wardlaw, Dieter Egli, Deyou Zheng, Claudia A. Doege, Rudolph L. Leibel

## Abstract

Bardet-Biedl Syndrome (BBS) is a rare autosomal recessive disorder caused by mutations in genes encoding components of the primary cilium and characterized by hyperphagic obesity. We developed a cellular model of BBS using induced pluripotent stem cell (iPSCs)-derived hypothalamic arcuate-like neurons. BBS mutations *BBS1*^M390R^ and *BBS10*^C91fsX95^ did not affect neuron differentiation efficiency but caused morphological defects including impaired neurite outgrowth and longer primary cilia. Expression of intact *BBS10* normalized cilia length. Single-cell RNA sequencing (scRNA-seq) of BBS1^*M390R*^ hypothalamic neurons identified several down regulated pathways including insulin and cAMP signaling, and axon guidance. In agreement with scRNA-seq data, insulin-induced AKT phosphorylation at Thr308 was reduced in *BBS1M390R* and *BBS10c91fsX95* human fibroblasts and iPSC-derived neurons, as well as in *BBS10* knockdown iPSC-derived neurons. Overexpression of intact *BBS10* fully restored insulin receptor tyrosine phosphorylation in *BBS10c91fsX95* neurons. Mutations in *BBS1* and *BBS10* impaired leptin-mediated p-STAT3 activation in both human primary fibroblasts and iPSC-derived hypothalamic neurons. Correction of the BBS mutation by CRISPR rescued leptin signaling. *POMC* expression in *BBS1*^M390R^ and *BBS10* ^C91fsX95^ iPSC-derived hypothalamic neurons was downregulated, as was hypothalamic *Pomc* in BBS1^M390R^ knockin (KI) mice. In the aggregate, these data provide insights into the anatomic and functional mechanisms by which components of the BBsome in CNS primary cilia mediate effects on energy homeostasis.

## Introduction

Bardet-Biedl Syndrome is an autosomal recessive disorder characterized by severe, early-onset obesity, as well as an array of other diagnostic features typical of ciliopathies including polydactyly, retinal degeneration, renal cysts and intellectual impairment (1). To date, 21 genes have been implicated in specific instances of BBS, and sixteen have been confirmed as components of the basal body of the primary cilium (PC) (2). Eight BBS-associated genes, BBS1, 2, 4, 5, 7, 8, 9 and 18, encode proteins that form a complex denominated the “BBSome” (3). The BBSome mediates transport of molecular cargo to the base of the PC via the Golgi or recycling endosomes (4), as well as ciliary exit by directing intra-flagellar transport (IFT)-loaded cargo out of the PC (5, 6). Ciliary cargo includes the receptors for SHH, Wnt, PDGFα, and other signaling molecules that are essential for embryonic development, cell growth, cell migration and cell survival (7, 8). Proteins associated with the PC are also important for the regulation of energy homeostasis: in Alström syndrome autosomal recessive mutations in the *ALMS1* gene - encoding a ciliary basal body protein - also cause obesity (9). Loss-of-function mutations in adenylate cyclase 3 (*ADCY3*) or melanocortin 4 receptor (*MC4R*), both of which localize to neuronal primary cilia, cause severe obesity in humans (10–14). In mice, systemic congenital deletions of *Bbs* genes, *Alms1*, *Adcy3* or *Mc4r* recapitulate the obesity phenotype (15–21). Hypomorphism of ciliary genes *Ift88, Kif3a* and *Rpgrip1l* results in increased adiposity in mice (22–24). Deletions of BBS-associated genes also cause obesity in mice: *Bbs1*^M390R/M390R^, *Bbs2^−/−^, Bbs4 ^−/−^, Bbs6 ^−/−^ and Bbs7 ^−/−^* mice are hyperphagic and obese (17–20).

The mechanisms by which BBS proteins affect body weight regulation are not well understood. BBS2 or BBS4 deletion in mice is associated with failure of G protein coupled receptors (GPCRs) that regulate energy balance such as somatostatin receptor 3 (SSTR3) and melanin concentrating hormone receptor 1 (MCHR1) to localize to the PC in the central nervous system (25). Similarly, BBS1 deletion in mice prevents localization of the neuropeptide Y 2 receptor (NPY2R) to the PC in the arcuate hypothalamus; and BBS1 has been implicated in trafficking of the obesity-implicated serotonin 2C receptor (5-HT2CR) in the PC (26). BBS1, a BBSome structural protein that facilitates recruitment of membrane-bound proteins to the PC (3), also interacts with isoform b of the leptin receptor (LEPRB), which binds RPGRIP1L at the transition zone of the PC (23, 27–29). The *BBS1* mutation *M390R* prevents the BBS1-LEPRB interaction, suggesting that BBS1 may participate in leptin signaling by influencing leptin receptor trafficking to the vicinity of the PC (27). In mouse embryonic fibroblasts in which *Bbs1* is “knocked-down”, or in human primary fibroblasts homozygous for *BBS1*^M390R^, localization of the insulin receptor (IR) to the cell membrane is reduced (30). Consistent with these findings, pre-obese *Bbs2*^−/−^, *Bbs4*^−/−^ and *Bbs6*^−/−^ mice are hyperinsulinemic (30).

BBS broadly impacts the development of the central nervous system: MRI scans of BBS patients show distinct tissue- and region-specific brain abnormalities including reduced white matter in most brain regions, and reduced gray matter in the caudate/putamen (dorsal striatum), and hypoplasia of the hypothalamus and thalamus (31). Cells within the hypothalamus play a critical role in the control of body weight and systemic glucose homeostasis (32). In the arcuate hypothalamus (ARH), proopiomelanocortin (POMC)-expressing and Neuropeptide Y (NPY)/ Agouti related protein (AgRP)-coexpressing neurons “sense” peripheral hormones such as leptin, insulin, and ghrelin, and inhibit or increase food intake and energy expenditure (33). Conditional ablation of ciliary proteins encoded by *Tg737*, *Kif3a* or *Rpgrip1l* in POMC neurons results in obesity (22, 34). In mice, congenital ablation of *Bbs1* in POMC or AgRP neurons also increases adiposity (26). In *Bbs1*^M390R/M390^ mice, cilia of ependymal cells lining the third ventricle at the ARH are elongated with swollen distal tips, suggesting that BBS1 is also important for maintaining normal ciliary structure in the hypothalamus (20).

Based on the studies enumerated above, it is likely that ciliary function is deranged in BBS patient hypothalamic cells. The ability to generate brain region-specific neurons from patient-specific iPSC now enables analysis of molecular and structural phenotypes in processes important to the control of food intake and energy expenditure (35). And, scRNA-seq has been widely used in developmental biology, oncology, immunology and metabolic diseases research (36–40) to identify cell-type-specific regulatory relationships among genes (37–39). It is particularly useful for resolving cellular heterogeneity in human iPSC-derived functional cell types. Here, we use these technologies to examine specific aspects of the molecular neurobiology of obesity in human BBS.

## Results

### Generation of BBS and control iPSCs

Two BBS fibroblast lines (GM05948 and GM05950) were obtained from Coriell. Ten skin biopsies (for the establishment of fibroblast lines) were obtained from BBS and healthy control subjects attending the Naomi Berrie Diabetes Center of the Columbia University Irving Medical Center under IRB protocol (**Table S1**). The BBS patients were ascertained primarily by Dr. Tsang to whom they had been referred for evaluation of associated retinopathy. Their clinical phenotypes were consistent with the diagnosis of BBS and included obesity, retinitis pigmentosa, polydactyly, anosmia, renal cysts and intellectual impairment (**Figure S1A, B**). Clinical diagnoses were confirmed by Asper Ophthalmics genetic testing or whole exome sequencing, which were further validated by dideoxy-sequencing (**Figure S1C**). The genotype of each subject is indicated in **Table S1**. Fibroblasts derived from skin biopies from BBS and control subjects were reprogrammed into iPSCs using retroviruses expressing *OCT4, SOX2, KLF4, c-Myc* (41). The studies reported here used the following lines: BBS1A (iPSC derived from patient 1085 with a homozygous *BBS1 M390R* mutation); BBS1B (iPSC derived from patient 1097 with a homozygous *BBS1 M390R* mutation); BBS10A (iPSC derived from the GM05948 primary fibroblast line with a homozygous mutation in *BBS10* (*C91fsX95*)); BBS10B (iPSC derived from GM05950 fibroblasts segregating for a compound heterozygous mutations in *BBS10* (*S303RfsX3*) (**Figure S1C**), as well as Control1 and Control2. The generation of iPSC lines BBS1A, BBS1B, BBS10B, Control1 and Control2 was previously published (35), and BBS10A is described here. Immunocytochemistry (ICC) analysis revealed that BBS10A expressed pluripotency markers (**Figure S2A**). Teratomas generated by BBS10A iPSC included all three germ layers. BBS10A had a normal karyotype (**Figure S2B-C**). Both control and BBS iPSC lines expressed endogenous pluripotency markers and showed complete silencing of exogenous genes (viral transgenes) (**Figure S3**). An isogenic control iPSC line for BBS1B was generated using CRISPR. In this line, the M390R mutations on both alleles were corrected from TOPO subcloning (**Figure S4A-B**). The corrected BBS1B isogenic control line (referred to as c-BBS1B) had a normal karyotype and was pluripotent (**Figure S4C-E**).

### BBS mutations increase ciliary length in iPSC-derived TUJ1+neurons

Elongated, swollen PC on ependymal cells by the third ventricle have been observed in *Bbs1*^M390R/M390R^, *Bbs2*^−/−^, *Bbs4*^−/−^ and *Bbs6*^−/−^ mice, indicating that BBS proteins are essential for maintaining normal ciliary structure (20, 42). Here, AC3, ARL13B and Gamma-Tubulin were employed as markers in ICC analysis to determine ciliary length. In human fibroblast cultures, 40-50% of cells were ciliated (**Figure S5A**) and there was no difference among control, *BBS1* and *BBS10* mutant cells in the percentage of ciliated cells or ciliary length (**Figure S5 B-C**).

Both control and BBS iPSCs were converted into neurons using dual SMAD inhibition (**Figure S6A**) (43, 44). At day 30, more than 80% cells were TUJ1+, and there was no difference in percentages of TUJ1+ neurons among the control and BBS iPSC lines (**Figure S6B**). Control, *BBS1* and *BBS10* mutant iPSC-derived neurons were all ciliated (**Figure 1A**), but cilia in BBS mutant neurons were about 14% to 28% longer than those in control iPSC-derived neurons, especially in *BBS10* mutant neurons (**Figure 1B**). Whole cell current clamps revealed that control, *BBS1* and *BBS10* iPSC-derived neurons generated comparable action potential post current stimulation (**Figure S6C**). Biallelic mutations in *RPGRIP1L* (a transition zone protein) reduce ciliogenesis in both fibroblasts and iPSC-derived neurons (23, 34). Consistently, the expression of ciliary markers *ADCY3* and *IFT88* were reduced in the JBST (*RPGRIP1L* mutation) iPSC-derived neurons (**Figure 1C**). In contrast, mRNA levels of ciliary genes *ADCY3* and *ARL13B* were increased in BBS10A mutant cells (**Figure 1C**). Exposure to SHH/FGF8 resulted in further lengthening of the PC in BBS10A mutant neurons by 50%, but not in *BBS1* mutant or control neurons, suggesting divergent roles of *BBS1* and *BBS10* mutations on SHH/FGF8 signaling (**Figure S7**).

**Figure 1.**
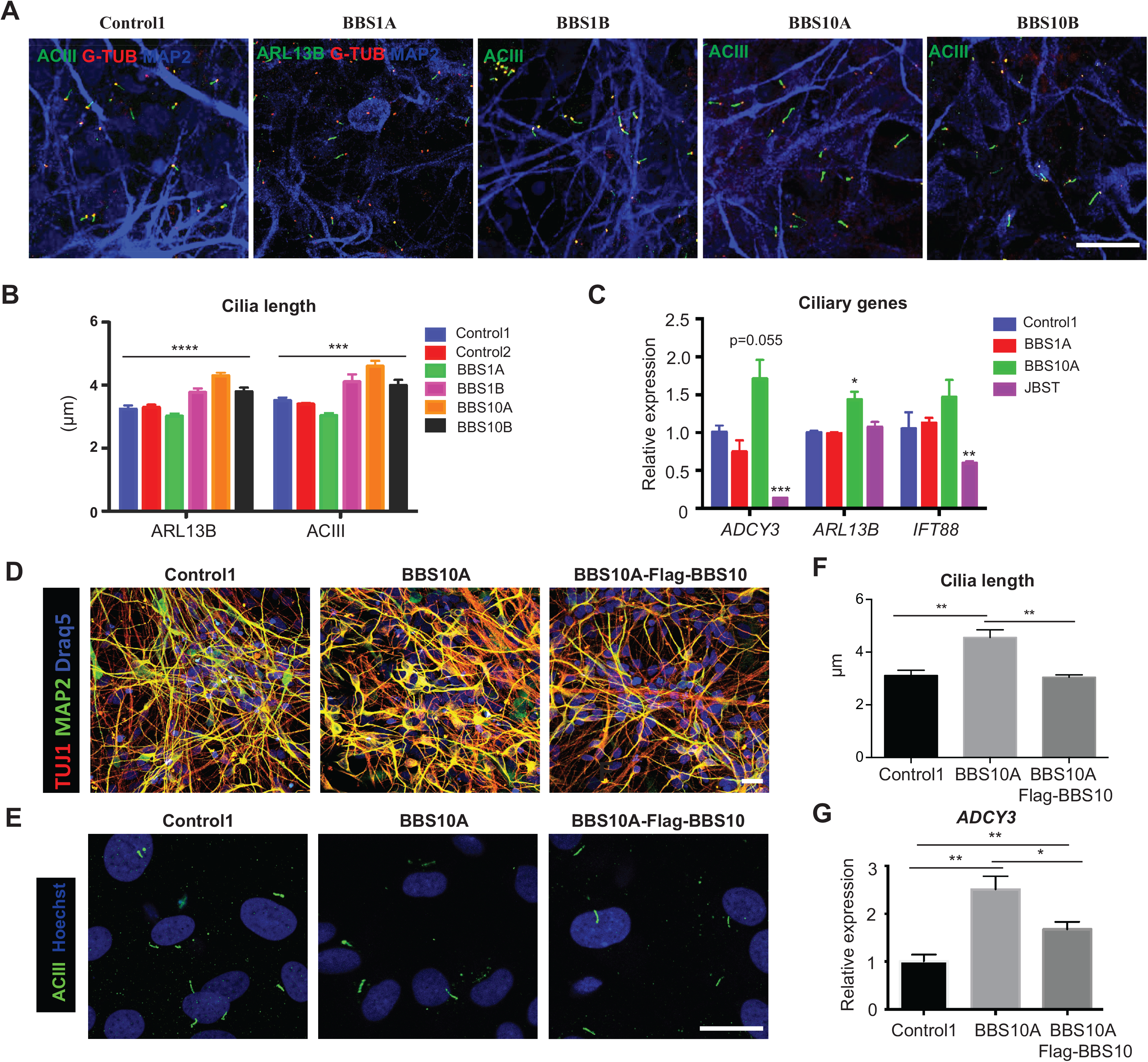
BBS mutations increase cilia length in iPSC-derived neurons. (A) Immunocytochemistry (ICC) staining of primary cilia in iPSC-derived neurons. Neurons were stained with MAP2 as well as cilia markers: adenylate cyclase III (ACIII, basal body and axoneme), ARL13B (axoneme) and G-TUB (basal body). Scale bar, 20μm. (B) Quantification of ciliary length in control and BBS iPSC-derived neurons. *** p<0.001, **** p<0.0001, One-way ANOVA analysis. (C) Expression of ciliary genes in control and BBS iPSC-derived neurons; Joubert syndrome (JBST) iPSC-derived neurons were used as control for impaired ciliogenesis. (D) ICC staining of *Control1*, *BBS10A* and *BBS10A-Flag-BBS10* iPSC-derived neurons. TUJ1 and MAP2 are neuronal markers. Scale bar, 20μm. (E) ICC staining of primary cilia in flow cytometry-sorted NCAM+ neurons. Cells were stained with ACIII and Hoechst. Scale bar, 10μm. (F) Quantification of cilia length. (G) Ectopic expression of Flag-BBS10 partially reverses *ADCY3* expression in BBS10A iPSC-derived neurons. Data are presented as mean±SEM, * p<0.05, ** p<0.01,*** p<0.001, student t-test.

In order to confirm that increased ciliary length was due to the BBS hypomorphic mutations, Flag-BBS10 was overexpressed in the BBS10A mutant line using lentivirus (**Figure S8**). Overexpression of Flag-BBS10 did not affect neuronal differentiation compared with control and BBS10A mutant lines (**Figure 1D**). AC3 staining of FACS-sorted NCAM+ neurons showed that overexpression of intact BBS10 normalized cilia length in BBS10 mutant neurons (**Figure 1E-F**). Gene expression analysis revealed that overexpression of Flag-BBS10 restored *BBS10* expression in the BBS10 mutant line and restored *ADCY3* expression back to normal (**Figure 1G, S8D)**. These data indicate that the BBS10 mutation elongates cilia and increases *ADCY3* in iPSC-derived neurons.

### Molecular signatures of *BBS1* mutant and its isogenic control hypothalamic neurons as obtained by scRNA-seq

BBS1B and c-BBS1B hypothalamic ARH-like neuron cultures were generated as previously described (35) and subjected to scRNA-seq using the 10X Genomics Chromium platform. For each line, we obtained more than 3,400 valid cells (>= 200 expressed genes) after filtering, with over 21,000 genes detected in >=3 cells (**Figure 2A**). Data from BBS1B and c-BBS1B iPSC-derived hypothalamic neurons were integrated and sorted into 14 clusters based on their signature genes (**Figure 2B, S9A, Table S5**). Clusters 1-4 and 7-10 share very similar gene expression profiles and were considered as differentiated neurons based on the presence of neuronal markers (including TUBB3, SNAP25, TH, GAD1, GAD2) (**Figure 2C**). POMC was highly expressed in clusters 3 and 10; SST was expressed in clusters 4 and 7 (**Figure 2C, S9B**). TRH was expressed in clusters 1, 2 and 8. The remaining clusters (expressing MAP2 but not TUBB3) show characteristics of neuron progenitors (**Figure 2B, C**). In particular, clusters 6 and 11-14 were enriched for neuron progenitor marker SOX2 (**Figure 2C**). Stem cell, astrocyte or oligodendrocyte markers such as NANOG, GFAP, ALDH1L1, S100B, OLIG1 and PDGFRA were either absent or expressed at very low levels (**Figure 2C**). Overall, the scRNA-seq analysis confirms hypothalamic neuron lineage and the anticipated heterogeneity of the human iPSC-derived ARC-like hypothalamic neurons.

**Figure 2.**
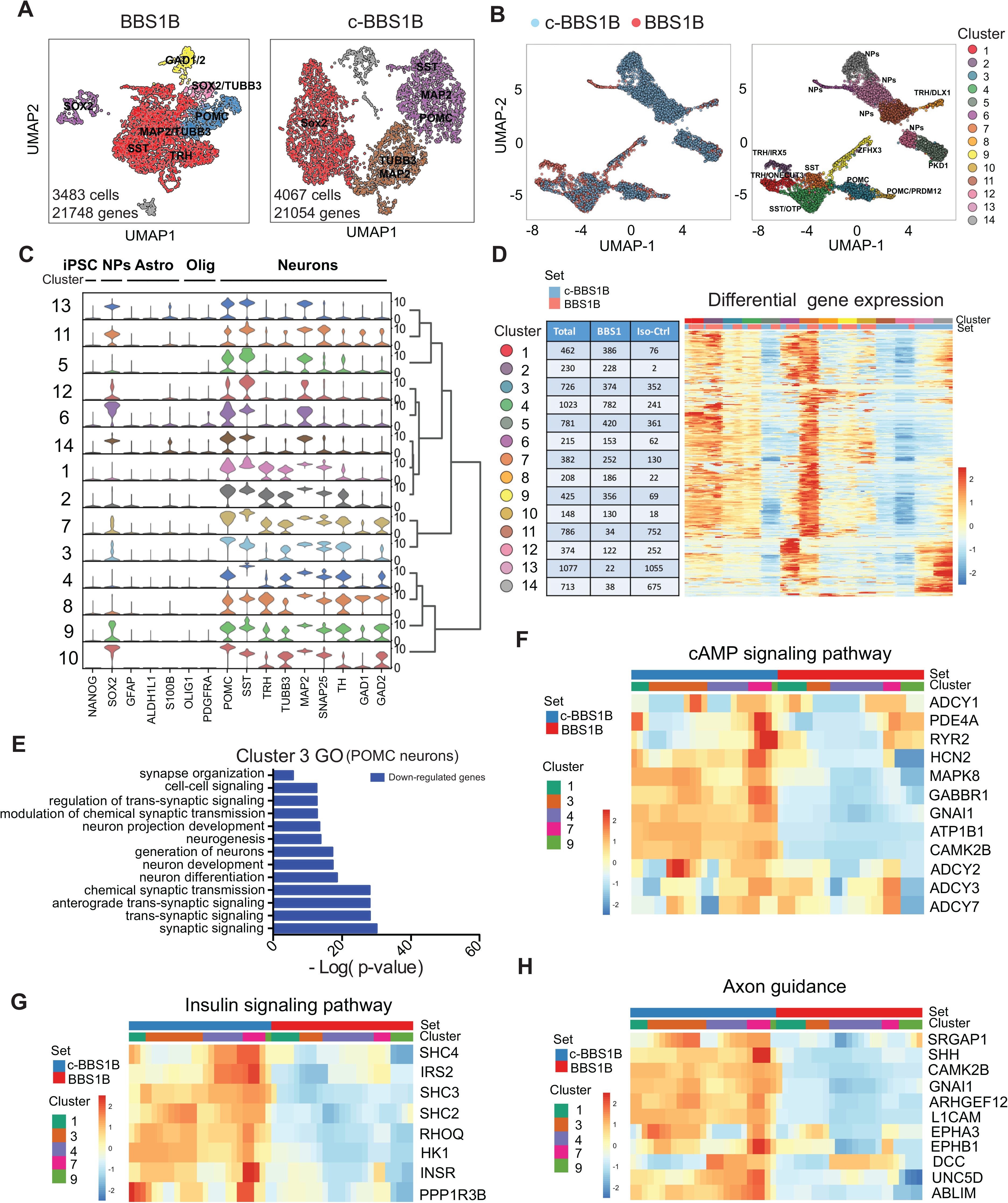
scRNA-seq analysis of BBS iPSC-derived hypothalamic neurons. (A) UMAP of *BBS1B* and isogenic control *c-BBS1B* iPSC-derived hypothalamic neurons. (B) UMAP of *BBS1B* and *c-BBS1B* neurons after integration and identification of 14 clusters. Marker genes and cell types for each cluster were indicated. (C) Violin plots of cell type-specific markers for all 14 clusters and hierarchical-clustering. Signature genes for iPSC, neuron progenitors (NPs), astrocytes (Astro), oligodendrocytes (Olig) and neurons were included. (D) Heatmap of differentially expressed genes in all clusters. No. of cells from which line within each cluster has been indicated in the table. (E) Gene Ontology (GO) analysis of down-regulated gene (BBS1B/c-BBSB) identified from Cluster 3. (F-H) Heatmaps of genes of pathways highlighted in KEGG pathway analysis. Genes involved in cAMP signaling pathway (F), insulin signaling pathway (G) and Axon guidance (H) were plotted against cell set and cluster ID.

Next we performed differential expression analysis of each cluster between the BBS1B mutant and isogenic control c-BBS1B lines to explore cell type-specific effects of BBS hypomorphism (**Table S6**). As shown in the differential gene expression heatmap, all clusters contained differentially expressed (by BBS1 genotype) genes except cluster 2 that contained only a couple of cells from c-BBS1B line (**Figure 2D, Figure S9A**). Gene Ontology (GO) and KEGG pathway analysis were performed for genes that were either significantly up- or down-regulated in BBS1B vs c-BBS1B neurons. Specifically, GO analysis of down-regulated genes in cluster 3 revealed that synaptic organization, trans-synaptic and synaptic signaling, chemical synaptic transmission, neurogenesis and neuron development were negatively impacted in the BBS1B mutant POMC+ neurons (**Figure 2E**). Neuron projection development and synaptic functions were also negatively impacted in BBS1B mutant cells of the other neuronal clusters 1, 4 and 7-10 (**Figure S10**). KEGG pathway analysis (**Figure S11**) revealed that transcripts related to cAMP signaling (**Figure 2F**), insulin signaling (**Figure 2G**), axon guidance (**Figure 2H**) and Type II diabetes mellitus (**Figure S12**) were diminished in BBS1B mutant neurons in clusters 1, 3, 7 and 9.

### BBS mutations impact neurite outgrowth and diminish Wnt and SHH signaling in iPSC-derived neurons

Neuroanatomic alterations in BBS patients include ventriculomegaly, thinning of the cerebral cortex and reduction in size of the hippocampus and corpus striatum (31). Based on the gene expression results of the iPSC-derived hypothalamic neurons described above, we investigated the impact of BBS hypomorphism on neurogenesis in iPSC-derived TUJ1+ neurons. Since BBS mutations did not affect the differentiation efficiency of iPSCs into TUJ1+ neurons we investigated the numbers of processes and lengths of neurites two days after re-plating of day 30 control and BBS iPSC-derived neurons. *BBS1* and *BBS10* iPSC-derived neurons showed reductions in both number of processes and neurite length (**Figure 3A-C).**

**Figure 3.**
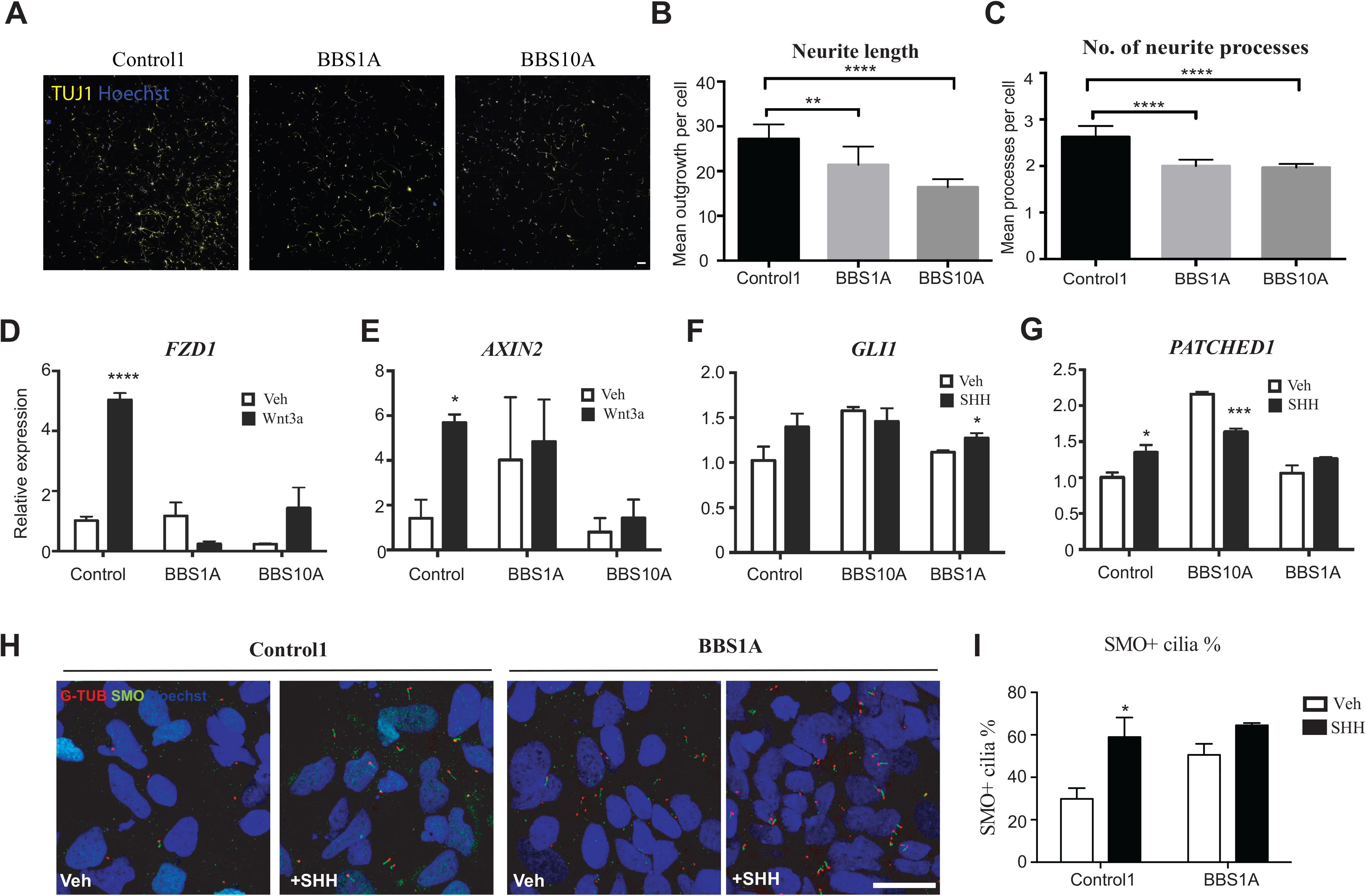
BBS mutations disturb neurite outgrowth and impair Wnt and SHH signaling in TUJ1+ iPSC-derived neurons. (A-C) Neurite outgrowth assay of Control1, BBS1A and BBS10A iPSC-derived neurons at day 30 of differentiation. (A) TUJ1 staining of Control and BBS mutant cultures. Scale bar, 50μm. Mean neurite length (B) and average number of neurite processes (C) were calculated using neurite outgrowth tool in metamorph software based on TUJ1 and Hoechst staining. (D-E) Wnt signaling is impaired in BBS iPSC-derived neurons. Control1, *BBS1A* and *BBS10A* iPSC-derived neurons (Day 30) were treated with Vehicle (Veh) or 100 ng/ml Wnt3a for 16hrs. Frizzled 1 (*FZD1)* and *AXIN2* mRNA levels were determined by Quantitative PCR (QPCR). (F-G) SHH signaling is reduced in BBS iPSC-derived neurons. Control and BBS iPSC-derived neurons (Day 30) were treated with Veh or 100ng/ml SHH for 16hrs. *GLI1* and *PATCHED1* mRNA levels were analyzed by QPCR. (H) Smoothened (SMO) staining in SHH-treated control and BBS iPSC-derived neurons. Control and BBS1A iPSC-derived neurons were treated with Veh or100 ng/ml SHH overnight. Neurons were fixed and stained with SMO, ϒ-Tubulin (G-TUB, basal body) for cilia and Hoechst for nuclei. Scale bar, 20μm. (I) Quantification of SMO+ cilia in E. Hoechst was used as nuclear marker. * p<0.05, ** p<0.01, *** p<0.001, **** p<0.0001 by 2-tailed student t-test.

Wnt and SHH signaling are critical for neurogenesis and neuron migration during brain development (34, 45–48). The trafficking and localization of SHH receptors, Smoothened (SMO) and Patched, is tightly regulated at the PC, and Wnt signaling is activated near the base of the PC (49). Wnt and SHH signaling were further investigated in TUJ1+ neurons. Wnt3a exposure strongly induced the expression of Wnt downstream targets *FZD1* and *AXIN2* in control neurons but not in BBS mutant lines (**Figure 3D, E**). Similarly, SHH exposure of control neurons increased the expression of its target genes *GLI1* and *PATCHED1*, a response that was strikingly reduced in BBS10A and decreased in BBS1A mutant neurons (**Figure 3F, G**). The number of SMO-positive cilia following SHH induction was significantly reduced in BBS mutant compared with control iPSC-derived neurons (**Figure 3H, I**). Blunted neurite outgrowth and defective Wnt and SHH signaling further implicate BBS proteins in important aspects of neuronal development.

### BBS mutations impair insulin signaling through interactions with the insulin receptor

Insulin signaling mediates aspects of energy and glucose homeostasis centrally and peripherally (50, 51). In the hypothalamus, insulin acts in a manner similar to leptin to reduce food intake by activating POMC neurons and inhibiting NPY/AGRP neurons (50, 52). Deletion of *Bbs2*, 4 or 6 results in hyperinsulinemia in mice independent of obesity, and the *M390R* homozygous mutation in *BBS1* fibroblasts results in perturbed IR trafficking to the cell membrane (30). It has been reported that IGF1 receptor function may be PC-dependent in 3T3-L1 pre-adipocytes (53). To determine whether BBS proteins participate insulin signaling in humans, we assessed insulin sensitivity by quantifying AKT phosphorylation (p-AKT) in response to insulin in BBS fibroblasts. By western blotting, insulin-induced AKT phosphorylation at Thr308 was reduced by 50% and 80-90% in BBS1 and BBS10 fibroblasts, respectively, compared with fibroblasts from control subjects (**Figure 4A**).

**Figure 4.**
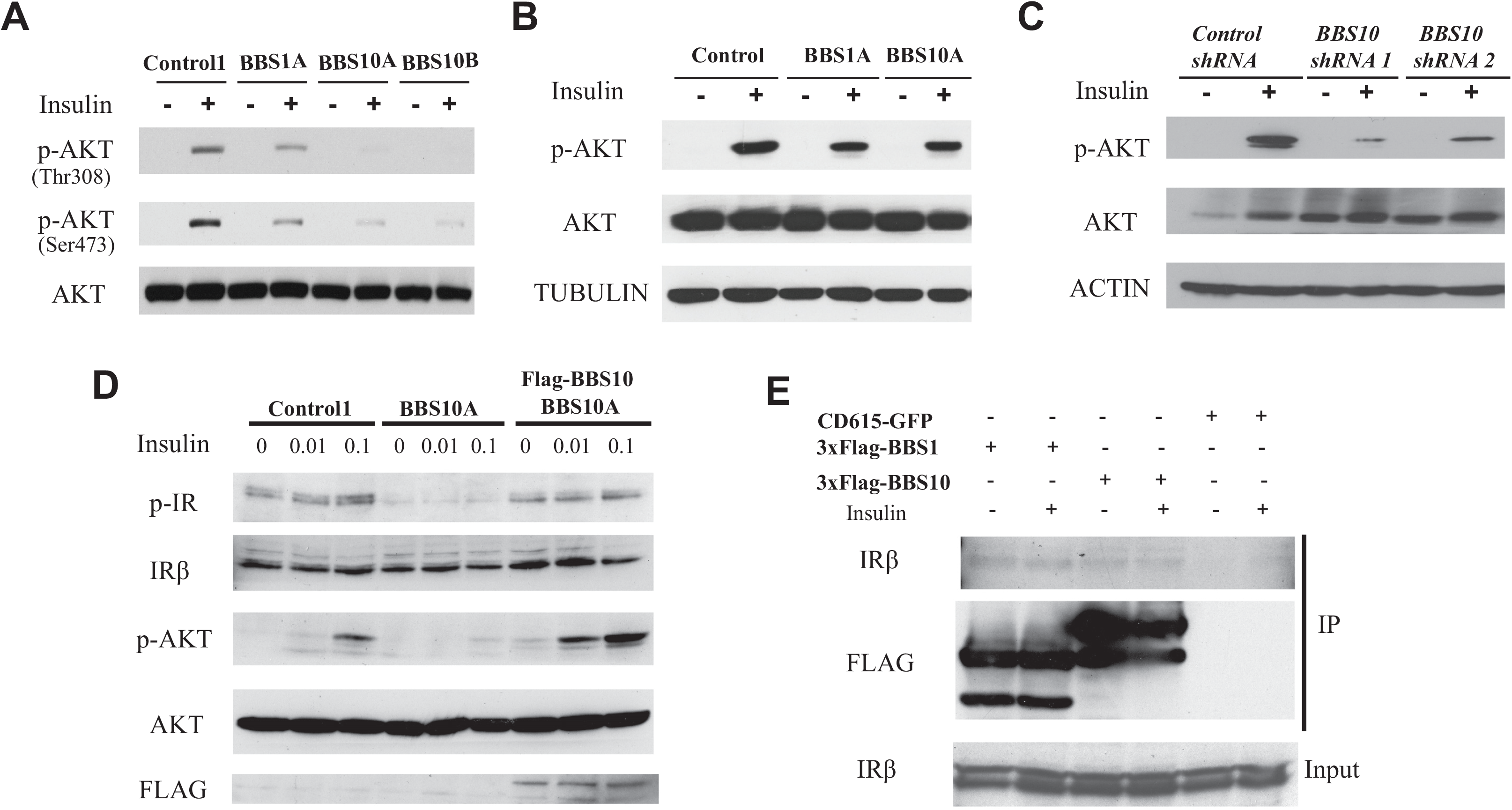
BBS1 and BBS10 bind to the insulin receptor and influence insulin signaling. (A) BBS mutations disrupt insulin signaling in human fibroblasts. Western blot (WB) analysis of insulin signaling as indicated by phosphorylation of AKT in control and BBS fibroblasts. Fibroblasts were serum-starved for overnight and treated with 0 or 1 μg/ml insulin for 30min. Both p-AKT Thr308 and p-AKT Ser 473 and total AKT were probed. (B) BBS mutations abrogate insulin signaling in iPSC-derived TUJ1+ neurons. Control, *BBS1A* and *BBS10A* iPSC-derived neurons (Day30) were fasted overnight and treated with 0 or 1 μg/ml insulin for 30min.p-AKT Ser 473 and AKT were probed. TUBULIN was used as a loading control. (C) Diminished insulin signaling in neurons derived from two *BBS10 shRNA* knockdown iPSC lines as indicated. iPSC-derived TUJ1+ neurons (Day30) were fasted overnight and treated with 0 or 1μg/ml insulin for 30min. Protein was collected for WB analysis of p-AKT Ser473, AKT and ACTIN. (D) BBS10 disrupts insulin signaling by disturbing phosphorylation of insulin receptor (IR) in Day12 neuron progenitors (NPs). WB analysis of insulin signaling molecules (p-AKT Ser473, AKT, p-IRβ and IRβ) in control, BBS10A and BBS10A-FLAG-BBS10 transgenic NPs after 30min treatment with 0.01 and 0.1 μg/ml insulin. FLAG was used to confirm the overexpression of wildtype BBS10. (E) Co-immuno-precipitation (IP) confirms the interaction between BBS proteins and IR. 293FT cells were transfected with CD615-GFP or CD615-3xFlag-BBS1-GFP or CD615-3xFlag-BBS10-GFP with Lipofectamine 2000. Cells were harvested 48hrs post transfection for Co-IP analysis. IRβ and FLAG were probed in Flag IP samples and total input.

In iPSC-derived TUJ1+ neurons, insulin-stimulated p-AKT levels were decreased by ~50% in neurons of both *BBS1* and *BBS10* mutant genotypes (**Figure 4B**). To assess the role of BBS proteins in insulin signaling, we knocked down (KD) *BBS10* expression in control iPSCs using lentiviral shRNA (**Figure S13**). In neurons generated from iPSCs where *BBS10* was KD by 40-60%, insulin-induced phosphorylation of AKT was reduced by 70-90% (**Figure 4C**). In the BBS10A mutant iPSC-derived neuron progenitors, insulin-induced p-AKT level was decreased 80-90% and fully rescued by overexpression of intact BBS10 (**Figure 4D**). Tyrosine phosphorylation of the insulin receptor (IR) was also reduced 60-70% in the BBS10 mutant neuron progenitors, a molecular phenotype that was fully rescued by overexpressing intact BBS10 (**Figure 4D)**. To determine whether BBS proteins directly interact with the insulin receptor, we performed co-immunoprecipitation in 293FT cells overexpressing transgenic Flag-BBS1 or Flag-BBS10. BBS1 or BBS10 were co-immunoprecipitated with IR, irrespective of the presence of ambient insulin (**Figure 4E**). Thus, BBS1 and BBS10 may regulate insulin signaling by direct protein-protein interaction with the IR.

### BBS mutations curtail leptin signaling in iPSC-derived hypothalamic neurons independent of obesity

It is challenging to parse defects in leptin signaling that are independent of the obesity exhibited in various Bbs knockout mice (17, 18, 20, 54). Human fibroblasts do not express LEPRb and therefore do not increase p-STAT3 level upon leptin exposure *in vitro* (**Figure S14A**). We induced ectopic expression of RFP-tagged LEPRb (RFP-LEPRb) in both control and BBS human fibroblasts using a lentivirus (**Figure S14B-C**). By Western blot analysis RFP-LEPRb was equally expressed in control and BBS fibroblast lines (**Figure 5A**). Leptin-induced p-STAT3 levels were reduced ~50% in leptin-stimulated RFP-LEPRb-transfected BBS1A fibroblasts, and completely blunted in transfected BBS1B and BBS10A fibroblasts compared to the control (**Figure 5A-B, S14D**). p-STAT3 immunostaining in leptin-treated transgenic fibroblasts revealed fewer leptin-induced p-STAT3 positive cells in BBS10A compared with control fibroblasts (**Figure S14E**), consistent with diminished leptin sensitivity of the BBS1B and BBS10A primary fibroblast lines. Overexpression of intact *BBS1* in BBS1B fibroblasts partially restored leptin signaling as reflected in partially restored p-STAT3 protein levels (**Figure 5B),** and increased p-STAT3 positive cells (**Figure 5C**). Leptin signaling was also assessed in iPSC-derived neurons which constitute an *in vitro* model without any influences of systemic adiposity. In TUJ1+ neurons at differentiation day 31, levels of *LEPRb* mRNA were comparable among control, BBS1A and BBS10A iPSC-derived neurons (**Figure S15A**). However, exposure of both control and *BBS* mutant neurons to leptin did not increase phosphorylation of STAT3 (data not shown), nor was expression of the leptin signaling negative feedback regulator *SOCS3* increased (**Figure S15B**), suggesting that the canonical leptin pathway is not operational in these neurons. Potential explanations include low expression of LEPRb and improper neuronal cell type in which leptin signaling is functionally inconsequential. To address this issue, RFP-LEPR was transiently overexpressed in control and *BBS* iPSC-derived Day 12 neuron progenitors (**Figure 5D**). In these cells, exposure to leptin increased p-STAT3 in the control but was weak or undetectable (**Figure 5D-F**) in the *BBS1* or *BBS10* mutant iPSC-derived neuron progenitors. In Day34 iPSC-derived hypothalamic neurons, we found that POMC-expressing isogenic control c-BBS1B and control neurons responded to leptin (p-STAT3 immunostaining), while both BBS1B and BBS10A mutant lines did not increase p-STAT3 in response to leptin exposure (**Figure 5G**). Collectively, these experiments suggest that BBS mutations contribute to diminished leptin sensing in the central nervous system independent of intercurrent increased adiposity.

**Figure 5.**
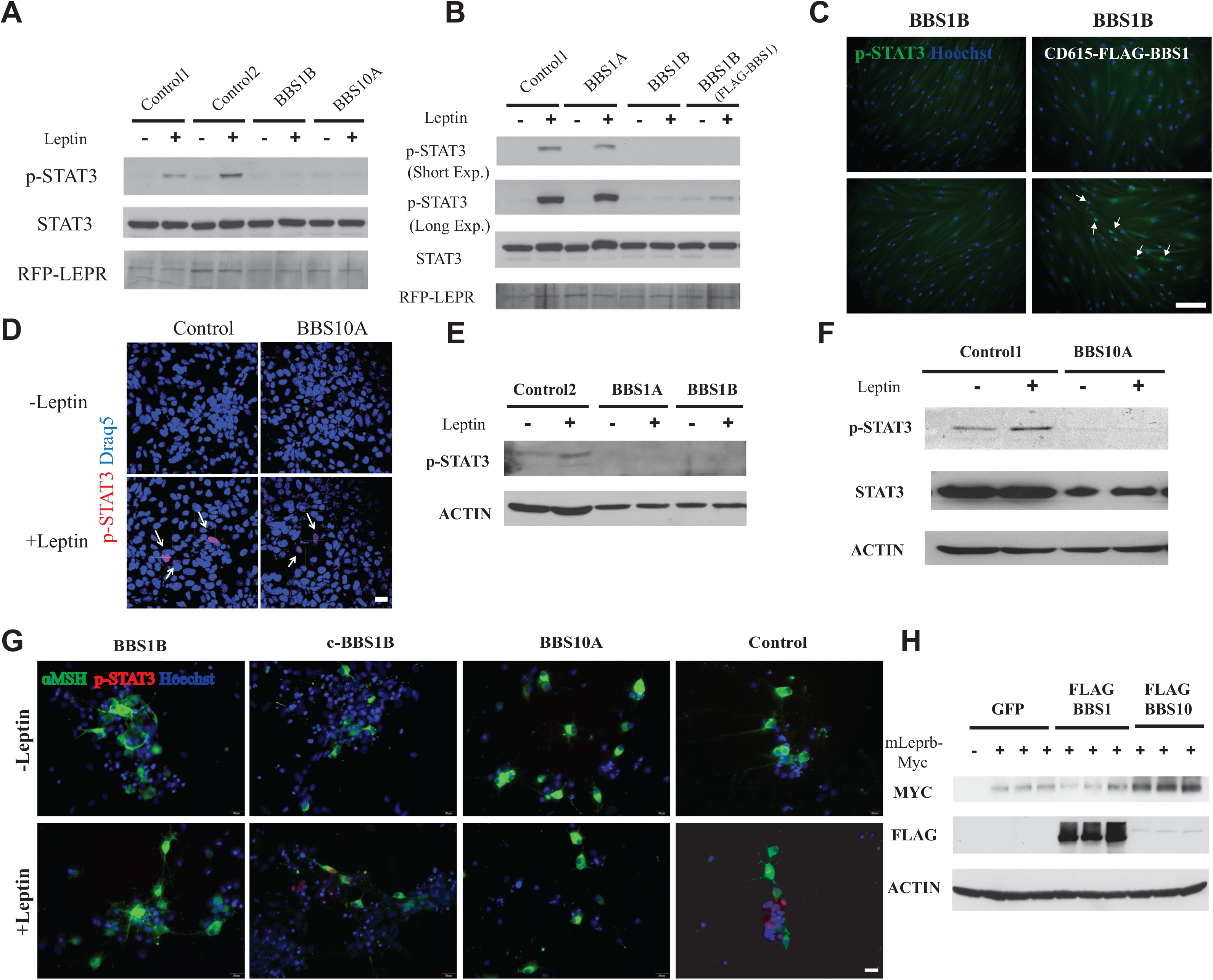
BBS mutations impair leptin signaling in human fibroblasts (RFP-LEPR transgenic) and iPSC-derived hypothalamic neurons. (A-B) Leptin signaling is impaired in RFP-LEPR transgenic BBS fibroblasts. Control, BBS1A, BBS1B and BBS1B (coexpressed Flag-BBS1) RFP-LEPR transgenic fibroblasts were treated with 0 or 0.5 μg/ml leptin treated (30min) after overnight serum starvation. Leptin signaling was measured by p-STAT3. STAT3 and RFP were also probed; Exp: exposure time. (C) Leptin signaling can be rescued in BBS1B RFP-LEPR transgenic fibroblasts by overexpressing wildtype BBS1. Leptin signaling was measured by the immunostaining of p-STAT3 in BBS1B and BBS1B+Flag-BBS1 RFP-LEPR transgenic human fibroblasts. Cells were fasted overnight and exposed to 0 or 1 μg/ml leptin for 30min followed staining with p-STAT3 and hoechst. Scale bar, 200μm. (D) Defective leptin signaling in BBS10A iPSC-derived neuron progenitors. Day12 Control and BBS10A iPSC-derived neuron progenitors were transfected with RFP-LEPR for 48 hrs. Cells were fasted overnight and then treated with 0 or 1ug/ml leptin for 30min. Immunostaining of p-STAT3 was performed. Scale bar, 20um. (E-F) Leptin signaling is disturbed in BBS iPSC-derived hypothalamic neurons. p-STAT3 was used as an indicator for leptin signaling. ACTIN and STAT3 were used as loading controls. (G) Leptin signaling as measured by immunostaing of p-STAT3 is disrupted in BBS iPSC-derived hypothalamic neurons. Day34 iPSC-derived hypothalamic neurons were serum fasted for overnight and treated with 0 or 1μg/ml leptin for 30min. Neurons were further stained with αMSH, p-STAT3 and Hoechst. Scale bar, 20μm; (H) Overexpression of *BBS10* increases total LEPR proteins. 293FT cells were co-transfected with mouse Leprb-Myc (mLeprb-Myc) and GFP or Flag-BBS1 or FLag-BBS10 for 24 hrs. MYC, FLAG and ACTIN were probed.

In order to further assess the role of BBS proteins in the regulation of leptin signaling, 293FT cells were co-transfected with mouse Leprb Myc-tagged in the C-terminus, and wild type human Flag-BBS1/BBS10, and assayed for the total amount of LEPRb protein by western analysis. Cells co-transfected with Flag-BBS10 contained 3-4-fold more LEPRb protein than GFP- or Flag-BBS1 co-transfected cells, suggesting that BBS10 enhances LEPRb translation and/or reduces protein degradation (**Figure 5H**). BBS10 proteins may participate in aspects of leptin signaling by effects on the stability of LEPRb. BBS1 mutations may impair leptin signaling by disrupting the trafficking of LEPR (27). Our findings suggest that BBS10 regulates leptin signaling by promoting molecular stability or trafficking of LEPR in iPSC-derived neurons.

### BBS1M390R mutation increases body weight and reduces hypothalamic POMC production

Previous reports claimed that hypothalamic *Pomc* mRNA level was decreased in several Bbs mice (19, 27). To further explore the potential effects of the BBS1M390R mutation on POMC expression, we studied BBS1M390R KI mice (20). On regular (6% fat) chow, KI mice displayed lower body weight during the first 6 weeks of age compared with littermate control mice (**Figure 6A**). At 10 weeks of age, the KI mice had the same body weight as control mice and performed comparably during the glucose tolerance test (**Figure 6B**). However, at 12 weeks of age the KI mice fed breeder chow (10% fat) had increased body weight and diminished glucose tolerance compared to control mice (**Figure 6C-E**). By 24 weeks of age, KI mice weighed 15% more than control littermates due to 40% and 10% increased fat and lean mass, respectively (**Figure 6F-G**). Fasting glucose in 24-week-old KI mice was higher than control mice (**Figure 6H**). However, the glucose tolerance of these KI mice was not affected (**Figure 6H**). Finally, in hypothalami of 25-week old KI mice, *Pomc* transcript was reduced by ~40% while *Npy* expression was unaffected (**Figure 6I**). The reduced hypothalamic Pomc expression in KI mice is consistent with studies of other Bbs mice (19, 27).

**Figure 6.**
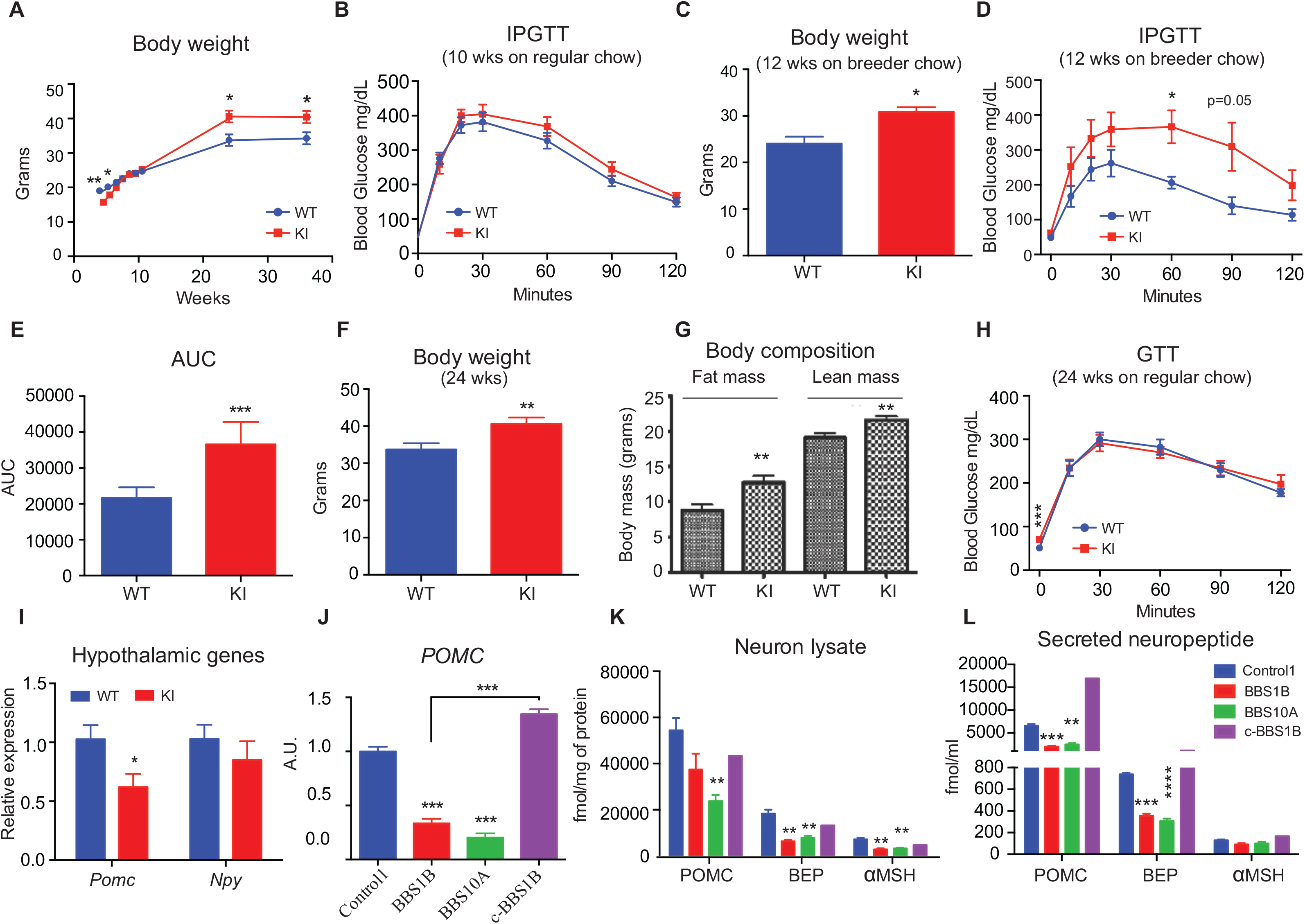
*BBS1M390R* mutation reduces *POMC* expression in both mouse hypothalamus and human iPSC-derived hypothalamic neurons. (A) Body weight curve of male WT and *BBS1M390R* knockin (KI) mice (n=9,10). (B) Intraperitoneal glucose tolerance test (IPGTT) of 10-wk old male mice on regular chow diet (n=9,10). (C-D) Body weight (C) and GTT (D) of 12-wk old male mice fed breeder chow *ad libitum* (n=6,7). (E) The glucose area under curve (AUC) in WT and KI mice as shown in D. (F) Body weight of 24-wk old WT and KI mice on regular chow diet (n=9,10). (G) Body composition of mice in F. Fat mass and lean mass were determined by TD-NMR. (H) GTT 24-wk old WT and KI mice on regular chow diet (n=9,10). (I) QPCR analysis of *Pomc* and *Npy* expression in mouse hypothalamus of WT and KI mice (24-wk old male) after 16hr fasting followed by 4hr refeeding (n=9,10). (J) QPCR analysis of *POMC* expression in Day35 iPSC-derived hypothalamic neurons. (K-L) Amount of neuropeptide produced in neuron lysates (J) and in cultured medium (16hrs culture, K) from control, BBS1B, BBS10A and c-BBS1B iPSC-derived hypothalamic neurons. POMC, αMSH and BEP concentrations were measured with ELISA and normalized to total protein. All comparisons were made to control iPSC-derived neurons. * p<0.05, ** p<0.01, *** p<0.001, **** p<0.0001 by multiple student t-test.

As previously reported, there were no gross differences in the percentage of NKX2.1+ and POMC+ hypothalamic neurons derived from control and BBS iPSC (35). *POMC* mRNA levels were reduced in BBS mutant iPSC-derived hypothalamic neurons (**Figure 6J**). BBS1B and BBS10A mutant human iPSC-derived hypothalamic neurons had lower amounts of alpha-melanocyte-stimulating hormone (αMSH) and β-endorphin (derivatives of POMC) in neuron lysates and in the growth medium compared to the control and c-BBS1B isogenic lines (**Figure 6K-L**), suggesting BBS proteins regulate POMC production.

## Discussion

We examined the molecular pathogenetic impact of the homozygous *BBS1* M390R and *BBS10* C91fsX95 mutations and the heterozygous *BBS10* S303RfsX3 mutation. Patients segregating for BBS10 mutations are generally more severely clinically affected than those segregating for BBS1 mutations (55). BBS10 is not a structural component of the BBSome but participates in the initial steps of the BBSome assembly by facilitating the interactions of BBSome structural components with canonical CCT chaperonins (54). These distinctions notwithstanding, we found that both BBS genes regulate ciliary structure, neurite anatomy, *POMC* expression, and responses to leptin and insulin. These findings are concordant with the anatomic and functional consequences of BBS mutations in human subjects with BBS1 or BBS10 mutations (31, 55).

In agreement with reports of abnormal ciliary structures in various BBS mouse models (20, 44), ciliary length in human stem cell-derived BBS mutant TUJ1+ neurons — and especially in *BBS10* mutant neurons — is significantly increased. Inhibition of AC3 activity promotes cilium lengthening in cultured synoviocytes (53). The up-regulation of *ADCY3* expression in the *BBS10* mutant neurons suggests that BBS10 may influence ciliary length via an AC3-mediated mechanism. scRNA-seq identified differential expression of genes involved in cAMP signaling pathway between BBS and its isogenic control neurons. Previous human and mouse genetic studies have implicated AC3 in body weight regulation (10–12). It is possible that BBS protein regulate body weight at least partially through an AC3-dependent cAMP signaling pathway.

The scRNA-seq analyses of BBS1B and c-BBS1B iPSC-derived neurons confirmed their hypothalamic lineage and revealed the heterogeneity of these differentiated cells as reflected by the identification of 14 cell clusters after integration both data sets. The hypothalamic ARH-like neuron differentiation protocol employed here yields predominantly POMC neurons, but also smaller fractions of other neuronal subtypes such as SST and TRH which have been previously detected in the mouse ARH using scRNA-seq (35, 56). Such heterogeneity of human pluripotent stem cell-derived neurons has been observed also by other investigators (38, 57). The application of scRNA-seq, however, enabled us to identify signaling and molecular pathways that were affected in specific neuron types.

Using ICC, no gross differences of neuronal differentiation efficiency were observed between BBS-mutant and control iPSC-derived TUJ1+ neurons. However, neurite outgrowth (number of mean processes and neurite length) was reduced in both *BBS1* and *BBS10* iPSC-derived neurons, suggesting a defective neurite outgrowth or disturbed neuron development in BBS lines. Wnt and SHH signaling pathways are important in the control of neuron projection development (58, 59). The trafficking and localization of SHH receptors Smoothened (SMO) and Patched is tightly regulated at the PC, whereas Wnt signaling is activated near the base of the PC (49). Thus, the neurite outgrowth phenotype may be resulting from the deficiencies in Wnt and SHH signaling pathways in *BBS* mutant cells as suggested by our studies. GO and KEGG pathway analysis of scRNA-seq data of BBS1B and its isogenic control line c-BBS1B highlight signaling related to axon guidance, neuron projection development and synaptic organization and transmission. In Drosophila, increased insulin signaling by knocking down *FoxO* and *PTEN* during metamorphic neuronal remodeling promotes neuronal growth and neurite branching (60). Moreover, leptin plays a neurotrophic role to promote neurite outgrowth of ARH neurons in neonatal mice (61). Therefore, the diminished insulin and leptin signaling in BBS mutant neurons shown in our cellular assays may contribute to reduced projection density.

Insulin signaling was impaired both in *BBS1* and *BBS10* mutant human fibroblasts, more severely so in *BBS10* cells. scRNA-seq data also indicated that expression of genes involved in the insulin signaling pathway were significantly reduced in BBS1B line compared with its isogenic control line. The differences in insulin sensitivity between the BBS1 and BBS10 mutant fibroblasts are consistent with the observed insulin resistance assessed by HOMA-IR in human subjects with BBS10 mutations. The severity of resistance in BBS10 patients is greater than that in BBS1 patients (55). BBS1 and BBS10 Co-IP with the insulin receptor (IR) and the BBS10 mutation decreases IR autophosphorylation in response to insulin. Thus, BBS1 and BBS10 are implicated in insulin signaling by a mechanism that involves direct interactions of BBS proteins with the IR that facilitate IR autophosphorylation.

Leptin signaling was reduced in *BBS1* and *BBS10* mutant human iPSC-derived hypothalamic neurons; genetic correction restored leptin signaling in these neurons. Because BBS1 and BBS10 are important for BBSome-associated ciliary trafficking, and the BBSome and transition zone of the PC have been implicated in LEPRB trafficking at the base of the cilium, it is plausible that BBS1 and BBS10 affect leptin sensitivity by effects on LEPRB trafficking to the PC (27, 54). In addition to LEPRB trafficking, BBS10 may enhance LEPRB protein stability and/or reduce its degradation, as overexpression of intact BBS10 in 293FT cells increased the amount of co-expressed HA-LEPR protein.

Decreased numbers of POMC neurons and levels of *Pomc* mRNA expression have been reported in hypothalami of *Bbs*2, *Bbs4* and *Bbs6* knockout mice, suggesting that *Bbs* mutations may affect neurogenesis and/or survival of POMC neurons (19, 27). In *Bbs1M390R* KI mice, hypothalamic *Pomc* mRNA was reduced compared with littermate controls. BBS1B and BBS10A mutant human iPSC-derived hypothalamic neurons were also characterized by reduced *POMC* mRNA expression as well as lower amounts of processed αMSH and β-endorphin in protein lysates and growth medium. These findings suggest that BBS proteins participate in the regulation of POMC production and processing.

In summary, we assessed the cell-molecular consequences of BBS mutations in patient-specific iPSC-derived hypothalamic neurons in an effort to identify mechanisms by which these molecules regulate ciliary processes related to weight homeostasis.

## Methods

### Research subjects and cell lines

All human subjects provided written informed consent prior to their participation in this study. Human subject research was reviewed and approved by the Columbia Stem Cell Committee and the Columbia IRB (Protocol IRB-AAAK6905). BBS patients were ascertained by Dr. Stephen Tsang, Department of Ophthalmology, Columbia University Irving Medical Center to whom they had been referred for evaluation of retinal phenotypes. The skin biopsies from these patients were obtained at the Naomi Berrie Diabetes Center. These lines are designated: Berrie Center skin biopsy-derived fibroblast lines 1085 (**BBS1A**) and 1097 (**BBS1B**), as well as the other 8 fibroblast lines (**Table S1**). They were screened for the two most prevalent mutations found in BBS subjects: *BBS1M390R* (c.1169T>G) and *BBS10C91fsX95* (c.271dupT) with PCR and Sanger sequencing. Both **BBS1A** and **BBS1B** are homozygous for the *BBS1 M390R* mutation (**Table S1**). The other two BBS fibroblasts lines were obtained from the NIGMS Human Genetic Cell Repository of Coriell Institute for Medical Research: GM05948 (**BBS10A**) and GM05950 (**BBS10B**) (**Table S1**). Genomic DNA was isolated from cultured fibroblast or blood with QIAmp DNA Mini kit (QIAGEN). The mutations in the two Coriell fibroblast lines — **BBS10A**, **BBS10B** – were identified by Asper Biotech and further confirmed by dideoxy-sequencing (Genewiz). See **Table S2** for genotyping primers.

### Animal studies

The BBS1M390R KI mice were obtained from Dr. Val Sheffield’s lab. Mice were maintained at room ambient 22° to 24°C with a 12-hour dark-light cycle (7am-7pm) in a pathogen-free barrier facility. Only male mice were included in this study. All mice were fed *ad libitum* regular chow diet (6% fat, Purina LabDiet 5053) or breeder chow diet (10% fat, Purina LabDiet 5058). Glucose tolerance tests were performed after 16hrs overnight fasting with intraperitoneal injection of 20% glucose (2g of glucose/kg body mass). Body weights were measured weekly. Body composition was determined by TD-NMR using an EchoMRI-100H body composition analyzer (EchoMRI, Houston USA). Mice were sacrificed after 16hr fast. NSG mice were purchased from The Jackson Laboratory (stock no. 005557) and fed regular chow ad libitum. All animal studies were approved by the Columbia IACUC.

### Generation and characterization of iPSC from human fibroblasts

Primary fibroblasts were converted into iPSC using retroviruses as described previously (35). All iPSCs lines were cultured on mouse embryonic fibroblast (MEF) cells in human ES (hESC) medium:500 ml knockout DMEM, 90 ml knockout serum, 6.5 ml GlutaMAX, 6.5 ml NEAA, 6.5 ml penicillin/streptomycin, 0.65 ml β-mercaptoethanol, and 10 ng/ml bFGF. Stable iPSC clones were characterized by immunostaining with pluripotency markers including NANOG and TRA-1-60 (35) and by qPCR analysis to confirm retroviral gene silencing and expression of endogenous pluripotency genes. Pluripotency was further confirmed by teratoma assay (35). BBS iPSCs were suspended as 5 million/200ul Matrigel (BD Biosciences) and subcutaneously injected into NSG mice (The Jackson laboratory, stock no. 005557). 2-3 months later, teratomas were isolated and fixed with 4% paraformaldehyde for further immunohistochemistry analysis. For karyotyping, control and BBS iPSC lines were cultured in T25 flasks according to Cell Line Genetics ® instructions and karyotyped by Cell Line Genetics ®. Reagents, if not otherwise specified, were all from Thermo Fisher Scientific.

### Generation of BBS isogenic control iPSC c-BBS1B using CRISPR-Cas9

BBS1 guide RNA (gRNA) was designed using the software from Zhang Feng’s lab at MIT (http://crispr.mit.edu). The sequences of the gRNA and single strand DNA (ssDNA) oligo were: BBS1M390RgRNA-CACCTCGAGTGGTCCTGATGAGG; BBS1ssDNA oligo-TTCCCCAACTAAACTCTGACGTCTCCACATAGGATGCAGTGACCAGCCTTTGCTTTGGCCGGTACGGGCGGGAGGACAACACACTCATCATGACCACTCGAGGTGAGTGGAGTCAGACCTGGCAAGGGCTTTGAAGTCGGGAGTGAAGGGACAGGCCTGCTTCTGGGGAAAGAGGAGGAG. The gRNA was cloned into the pGS-U6-gRNA plasmids by Genscript®. The pCas9_GFP plasmid was purchased from Addgene (#44719). After the BBS iPSCs became 90% confluent on 6-well MEF plate (2 wells), each well was transfected with Lipofectamine 3000 supplemented with 2.5μg pCas9_GFP plus 2.5μg pGS-U6-gRNA in 2ml hESC medium. 48-72hrs post transfection, GFP+ cells were sorted with fluorescence-activated cell sorting (FACS) and plated them onto 6-well MEF plate with hESC medium plus 10uM Rock inhibitor Y-27632 (Rocki, Selleckchem). Flow cytometry data were analyzed using the BD FACSDiva 8.0.1 software. After 7-14 days, individual hESC colonies were picked for clonal expansion and screened with Sanger sequencing (Genewiz) of the PCR products amplified around the targeted genomic loci and confirmed via TOPO cloning (TOPO™ TA Cloning™ Kit). The isogenic control line was further characterized for pluripotency and karyotype.

### Neuronal differentiation of iPSC

Human iPSC were cultured and maintained in hESC medium on mouse embryonic fibroblasts (MEF, 111,000 cells/well in 6-well plates) (35, 62). Upon confluence, iPSC cells were cultured for 4 days in EB medium plus dual SMADs inhibitors — 10μM SB43154 and 2.5μM LDN-193189 (Selleckchem) (35). For days 5-8, EB medium (hESC medium without bFGF) was replaced with N2 medium in steps - from 75% to 50% to 25% and to 0%, while maintaining SB and LDN at constant concentration (35, 43). Neuron progenitors were detached from plates on day 12 after incubating in Trysin LE incubator for 4 min at 37C. Neuron progenitors were washed twice with N2 medium and expanded for 2-3 passages in N2 medium plus SB/LDN on poly-L-ornithine and Laminin (PO/LA) coated plates(35). For further neuronal differentiation, cells were cultured on PO/LA coated plates at 50,000 cells per 24-well or 4-well plate, and at 500,000 cells per 6-well plate. After culturing 2-3 weeks in N2 medium supplemented with 10 μM DAPT (Selleckchem), B27 and 20 ng/ml BDNF (R&D), TUJ1+ differentiated neurons were obtained.

ARH-type hypothalamic neurons were differentiated using a previously described protocol based on dual SMADs inhibition plus SHH activation and inhibition of Notch signaling ((35, 62)). After Day 30, differentiated neurons were maintained in Brainphys neuronal medium (Stemcell technologies) for another 1-2 weeks for functional maturation.

### Plasmids and lentivirus production

Lentiviral cloning and expression vectors pCDH-UbC-MCS-EF1-Hygro (CD615B-1) and pCDH-EF1-MCS-T2A-Puro (CD520A-1) were purchased from System Biosciences Inc. (SBI). Human cDNA clones for LEPR transcript variant 1 (Catalog # HG10322-M), BBS1 (Catalog # HG10498-M) and BBS10 (Catalog # HG15095-G) were purchased from SBI. (63). Fluorescent expression vectors pEGFP-N3 (Clontech) and pCAG-DsRed (Addgene) were used in this study. Genes for GFP or RFP were cloned into the CD520/CD615 with BamHI and NotI digestion. CD520-RFP-LEPR was generated as described elsewhere (23). 3xFLAG-BBS1 cDNA was PCR-amplified from the BBS1 cDNA plasmid using the following primers: 5’-TCTAGA<u>TCTAGA</u>GCGAAGATGGACTACAAAGACCATGACGGTGATTATAAAGAT CATGACATCGATTACAAGGATCACGATGCCGCTGCGTCCTCATCGGA-3’; 5’-GGATCC<u>GGATCC</u>CAGCAGCTCAGGT CACAGGCGG-3’ and was further cloned into the CD615-GFP plasmid in the Xba I and Bam HI restriction sites to construct the CD615-3xFLAG-BBS1-GFP plasmid. Primers 5’ – ATA<u>TCTAGA</u>AGATATGGACTACAAAGACC ATGACGGTGATTATAAAGATCATGACATCGATTACAAGGATCACGATTTAAGTTCTATGGCCGCTGCAGG-3’ and 5’-primer: TTA<u>GCTAGC</u>CTTCTGATGTGATAGTTTATC TTCTG-3’ were used to amplify the 3xFLAG-BBS10 from the BBS10 ORF plasmid using CloneAmp HiFi PCR premix (Takara), which was further cloned into the CD520A-GFP plasmid via XbaI and Nhe I restriction sites. CD615-GFP and CD520-3xFLAG-BBS10-EGFP were digested with XbaI and BstBI to generate CD615-3xFLAG-BBS10-EGFP. Plasmid maxiprep amplification was performed using the QIAGEN Plasmid Maxi Kit (Catalog # 12162, QIAGEN). To prepare lentiviral stocks, HEK293 cells were cultured to 90% confluence in 10 cm diameter tissue culture dishes and transfected with 10 μg lentiviral expression vector (CD520-RFP-LEPR, CD615-3xFLAG-BBS1 or CD615-3xFLAG-BBS10) and 20 μg lentiviral packing vector mix with 45μl lipofectamine® 2000 in OptiMEM Reduced Serum Medium (Catalog #31985088). After 6 hours of incubation at 37°C, 5% CO_2_, the Opti-MEM Medium was replaced with the 12ml fibroblast medium: 500 ml DMEM medium containing 4.5 g/ml glucose (Catalog#11960044) and 50 ml FBS (Catalog#16000044) and 0.5-1 μg/ml Pen/streptomycin from a 100x stock solution. To remove cell debris from suspensions containing lentiviral particles, 48 hours after transfection, the supernatant containing the lentiviral particles was filtered through a 0.45 μm filter (Millipore).

### Generation of human transgenic cell lines

Primary human fibroblast lines were cultured and maintained in fibroblast medium. At confluence, 2 ml of the lentiviral supernatant harvested from HEK293 cells was added to each well in 6-well plates. After 2 days, the lentivirus supernatant was removed and fibroblast medium with 1 μg/ml puromycin or 50 μg/ml hygromycin was added until we obtained stable transgenic fibroblast lines that express RFP-LEPR or express both RFP-LEPR and CD615-3xFlag-BBS1-GFP. To construct human transgenic iPSC lines, 2ml of the lentiviral supernatant were added to each well on the 6-well cultured iPSCs (>90% confluent). After 48 hours, the lentiviral medium was replaced with hESC medium plus puromycin (1 μg/ml) or hygromycin (50-200 μg/ml, Life technologies) until stable iPSC lines that express CD615-3xFlag-BBS10-GFP were obtained.

### BBS10 Knockdown

BBS10 shRNA lentivirus particles were purchased from Sigma (St.Louis). The sequences of the two shRNAs selected were: CCGGCCTCAGAAAGTTCACAATCAACTCGAGTTGATTGTGAACTTTCTGAGGTTTTTTG (TRCN0000167181) and CCGGGCATTTATACCACACTCTATACTCGAGTATAGAGTGTGGTATAAATGCTTTTTTG (TRCN0000167794) (SHCLNV-NM_024685). The non-mammalian targeting shRNA (Cat # SHC002V, Sigma) was used as a negative control. iPSC cultures were established on 24-well MEF feeder cells. At 90% confluence, 10ul shRNA lentivirus particles were added to wells filled with 0.5ml hESC medium. After incubation for 48 hours with the shRNA lentivirus, the medium was removed and fresh hESC medium containing 1μg/ml puromycin was added for another 24-48 hours. New MEFs were added to each well to help the survival of the iPSCs. The selection step was repeated until pure puromycin resistant iPSCs were obtained.

### Neurite outgrowth assay

Day 30 iPSC-derived neurons were dissociated from plate after 5min Trypsin LE treatments and collected in N2/B27/Rocki medium. After centrifugation at 800 rpm for 5min, supernatant was removed, and the cell pellet was washed once with N2/B27 medium. Neurons were resuspended with 2ml N2/B27/Rocki medium and let cells go through the 5ml round bottom polystyrene tube with cell strainer cap (Fisher scientific) to make single cell suspension. Cells were counted with hemocytometer and 2500 neurons were added to each well on PO/LA coated 96-well plate (transparent bottom black plate). 48hrs later, neurons were fixed with 4% PFA for 10min and stained with TUJ1, Neurofilament antibodies (NF) and Hoechst. The whole picture of each well was taken by TROPHOS plate runner. Images were further analyzed by MetaMorph software to quantify neurite length, neuron body size, number of processes, and count the number of TUJ1-stained cells for each channel.

### Quantitative PCR Analysis

RNA was purified with the RNeasy Mini (QIAGEN) or plus micro kit (QIAGEN) according to the manufacturer’s instructions (QIAGEN, cat no). cDNA was generated with the GoScript Reverse Transcription kit (Promega) with 1μg RNA from each sample. 20 μl cDNA were diluted with 180 μl dH_2_O. GoTaq**®**qPCR kit (Promega) was used for qPCR analysis. See **Table S3** for qPCR primers.

### Western blot and Immunoprecipitation

Human fibroblasts or iPSCs were starved in 6-well plates or 10 cm tissue cultured dishes in serum free DMEM medium overnight. They were then treated with insulin (1 μg/ml) or leptin (0.5 μg/ml) or vehicle (PBS) for 30min at 37 °C, 5% CO2. After treatment, medium was removed, cells were rinsed once with cold PBS. For iPSC-derived neuron progenitors or neurons, we cultured cells in neurobasal™ medium supplemented with B27 minus insulin overnight. Variable doses of insulin or 1 μg/ml leptin, or vehicle for each, were then added to the neurobasal medium and incubation was allowed at 37 °C, 5% CO_2_ for 30 minutes. Neuron progenitors were cultured on PO/LA coated plate and transfected with the RFP-LEPR carrying plasmid using the TransIT-Neural transfection reagent (Mirus) for 48 hours before leptin treatment. After aspiration of the medium, cells were rinsed with cold PBS. 250ul of lysis buffer (20 mM Tris, pH 7.4, 150 mM NaCl, 2% Nonidet P-40, 1 mM EDTA, pH 8.0, 10% glycerol, 0.5% sodium deoxycholate, 0.2% semi-dehydroascorbate, supplemented with phosphatase and proteinase inhibitors, Catalog#78440) to each well to prepare protein lysates. 15 or 30 μg of proteins from fibroblasts, iPSCs, neuron progenitors or neurons were loaded to each lane on a 4-12% NuPAGE gel (Catalog#NP0335BOX). 293FT cells were co-transfected with Myc-Lepr(b) plamid (64) together with CD615-GFP, CD615-3xFlag-BBS1-GFP or CD615-3xFlag-BBS10-GFP with lipofectamine ® 2000. 40 μg of total protein from neuron samples or 293FT cell lysate were used. For immunoprecipitations, 293FT cells were transfected on a 10 cm tissue cultured dish with CD615-3xFlag-BBS1-GFP or CD615-3xFlag-BBS10-GFP or CD615-GFP with lipofectamine ® 2000. After 24 hours, 293 cells were switched into serum free DMEM medium overnight before insulin treatment. 1 μg/ml insulin were added to 293FT cells for 30 min at 37 °C, 5% CO_2_. Cells were rinsed with cold PBS before lysed cell with IP buffer (50 mM Tris pH7.9, 150 mM NaCl, 10% glycerol supplemented with protease inhibitors). For Co-immunoprecipitations (Co-IP), 1mg protein lysates were incubated with 30ul Flag antibody-conjugated agarose beads (Sigma) overnight at 4°C. Beads were washed four times with IP buffer and precipitates were eluted by boiling in non-reducing sample buffer (20 mM Tris, pH 7.4, 150 mM NaCl, 10% glycerol and 1% SDS). Information on all antibodies used in this study can be found in **Table S4**.

### Immunocytochemistry/Immunohistochemistry

Fibroblasts were cultured on 12mm diameter round glass coverslips (Neuvitro corporation) and fasted for 24hr to 48 hrs in serum free DMEM medium before fixation while iPSC-derived neuron were also cultured on PO/LA coated glass coverslips for further cilia imaging. Day34 iPSC-derived TUJ1+ neurons were dissociated with Trypsin LE and harvested into N2 medium. Cell pellet was further resuspended in 150ul PBS+0.5%BSA and stained with PE anti-human CD56 antibody (1:100, Biolegend) for 20min in dark. Cells without antibody staining were used as negative control. FACS-sorted CD56+ cells with BD Biosciences ARIA-IIu Cell Sorter and replated on PO/LA coated glasscoverslip as 2500 cells/well on 4-well plate for further cilia imaging. Cultured cells were fixed in 4% paraformaldehyde for 10min at room temperature. For p-STAT3 staining, cold methanol treatment is required for additional 3-5 min at 4 °C after fixation. The following staining procedure is the same as described previously (35). Teratomas were embedded in paraffin and cut into 8 μm thickness sections. These teratoma sections were stained with Hematoxylin and Eosin (H&E) (35). See **Table S4** for antibody information. Images were acquired with an Olympus IX71 epifluorescence microscope with Olympus DP30BW black and white digital camera for fluorescence, and DP72 digital color camera for H&E staining. Some images were acquired with a Zeiss LSM5 Pascal microscope. Cilia length was quantified based on the distance between the basal body and the tip of the axoneme (34).

### Neuropeptide assays

Neuropeptide assays on iPSC-derived hypothalamic neurons were performed as previously described (35). Briefly, all neurons were cultured in 12-well plates and initiated with 300,000 cells at day 12. For neuron lysate samples, medium was aspirated, and the differentiated neurons rinsed once with PBS, after which 250 μL of 0.1 N HCl was added to each well. Cells were harvested with a cell scraper and the cell lysates were transferred into 1.7 mL Eppendorf tubes. These samples were sonicated on ice with maximum power for 5 min. After centrifugation at 10,000 × *g* for 10 min at 4°C, the supernatants were collected for POMC, αMSH and βEP assays. Results are presented in femtomoles (fmol)/ml. Protein concentrations in neuron lysate were determined using a Pierce BCA protein assay kit. Neuropeptide concentrations were normalized to the amount of protein in each well for each cell line.

### Single-cell isolation, single-cell RNA sequencing and data analysis

At day 32 of differentiation, neurons were dissociated using the cell dissociation enzyme Accutase. Briefly, for each cell line, 1 ml of Accutase was added to one well of 6-well plate and incubated at 37 °C for 2 min. Next, the Accutase was carefully removed, replaced with 1 ml of BrainPhys Neuronal Medium supplemented with N2 Supplement-A (StemCell Technologies) and B27 supplement minus vitamin A (N2 / B27 BrainPhys Neuronal Medium), and immediately dissociated by gently pipetting (15-20 times with P1000) using low retention and wide orifice tips (Rainin). The cell suspension was transferred into a 50 ml Falcon ® tube. The well was rinsed with an additional 6 ml of N2 / B27 BrainPhys Neuronal Medium in order to collect remaining cells and were subsequently also added to the tube. The suspension was centrifugated for 5 min at 300 g at room temperature. After centrifugation, the pellet was resuspended in 1 ml of N2 / B27 BrainPhys Neuronal Medium by gentle pipetting (10 times with P1000) using low retention and wide orifice tips, and then transferred to a low bind microcentrifuge tube (Fisher Scientific). The suspension was then spined for 5 min at 300 g at 4 °C. After centrifugation, the pellet was resuspended in 100 or 200 μL of N2 / B27 BrainPhys Neuronal Medium initially by gentle pipetting with a P200 using low retention and wide orifice tips (Rainin) for 20-25 times followed by low retention and regular orifice tips (Rainin) for 10 times. Then, the suspension was diluted 1:2 with fresh medium and passed through a 40 μm cell strainer (Fisher Scientific) to obtain single cell suspension. Trypan Blue was used to determine number of viable cells. A viability between 60– 80% was obtained for all the samples. The single cell suspension for all the cell lines were kept on ice until use. Libraries were prepared using the Chromium Single Cell 3ʹ Reagent Kit v3 (10x Genomics Inc) and sequenced on an Illumina NovaSeq 6000.

Cell ranger (v3.1.0, 10x) was employed with default parameters to process the the scRNA-seq data (65). The human reference genome (GRCh38-3.0.0) was used for the alignment of reads obtained from10x Genomics. The Cell Ranger summaries indicated that both datasets passed the quality controls, with valid barcodes > 97%, mean reads per cell > 66,000, and median genes per cell > 3,700. The filtered gene-barcode matrix from the Cell Ranger outputs was used for subsequent analysis.

The raw counts of scRNA-seq data were filtered and normalized using the “RISC” package (v 1.0.0) (66). Genes expressed in less than 3 cells were discarded, and cells expressing < 200 genes were removed. At the end, ~4,000 valid cells were obtained for BBS1 iso-control(c-BBS1B), and ~3,500 valid cells for BBS1B M390R mutant, with >20,000 genes detected for each sample. The count normalization was performed independently for each sample, with counts log-transferred and weighted by sequencing depth. The batch effects between the two datasets were corrected using “RISC” package in R environment (v 3.6.3), by projecting all the cells from both datasets into a common reference space. The top principal components (PCs = 18) were used to build the reference space for the integrated data, containing 7550 valid cells and ~19,700 genes. We also used these PCs for cell clustering, based on Louvain method of “igraph” package (v1.2.5), with neighbors set to 20 (http://igraph.org). In total, 14 clusters were obtained for the integrated data. Cell types were identified according to the expression patterns of cell type marker genes that were expressed significantly higher in one cluster vs others. The “scanpy” package (v 1.5.1) was utilized to generate the stacked violin plot for cell type markers in Python environment (v 3.8.5)(67).

Differentially expressed (DE) genes of the integrated data were determined using a Negative Binomial generalized linear model in “RISC” package, at adjusted p-value < 0.05 and logFC (log2 fold change) >0.25 or <−0.25. Heatmaps of DE genes were generated using the R “pheatmap” package (v 1.0.12); the violin plots of DE genes were prepared by the Python “seaborn” package (v 0.10.1) (https://doi.org/10.5281/zenodo.883859). GO and KEGG pathway analysis were performed using the ToppGene Suite (https://toppgene.cchmc.org/enrichment.jsp), at the significance level of P value < 0.05.

### Electrophysiology study of iPSC-derived neurons

Patch clamp recording of iPSC-derived neurons was conducted as previously described (35).

### Statistics

Unless otherwise indicated, all graphical data are presented as mean ± SEM; significance was calculated using 2-tailed Student’s t tests. P value of less than 0.05 were considered significant.

## Author contributions

L. Wang, D. Egli and RL. Leibel designed this study; L. Wang performed the experiments, data analysis and generated the initial draft of the manuscript; CA. Doege and G. Stratigopoulos contributed to data analysis and manuscript composition. Y. Liu and D. Zheng performed the scRNA-seq data analysis and helped manuscript drafting; DJ. William performed the electrophysiology studies; R. Goland and SH. Tsang provided skin biopsies and blood cells from the BBS subjects; L. Sui and LC. Burnett helped with cell culture experiments. C. Leduc and L. Shang helped with mouse work. Y. Zhang provided the mouse Myc-ObRb-HA plasmid and suggestions on leptin signaling studies. M. De Rosa and CA. Doege helped with neuron differentiation and sample preparation for scRNA-seq; HJ. Glover contributed to the scRNA-seq data analysis.

## Acknowledgements

We thank Haiqing Hua and Aiqun Li for discussions related to this work; Yao Li and Jing Yang for help with collecting the biopsy samples; Val Sheffiled for BBSM390RKI mice; Kathryn Anderson (Memorial Sloan Kettering) for the Smoothened antibody; Tamara Caspary for the ARL13B antibodies; and L. Yang of the Diabetes and Endocrinology Research Center Pathology Core at Columbia University for histology analysis. Some of the studies reported here were conducted in the Genomics and High Throughput Screening Shared Resource at Columbia University (NIH/NCI P30CA013696). This research was supported by RO1 DK52431-23, the New York Stem Cell Foundation, a NYSTEM IIRP award (SDHDOH01-C32590GG-3450000), the Rudin Foundation, the Russell Berrie Foundation Program in Cellular Therapies, the Diabetes (P30 DK63608-16) and Obesity Research (P30 DK26687-41) Centers of Columbia University, RO1DK093920, U54OD020351, R24EY028758, U01 EY030580, R24EY027285, 5P30EY019007, R01EY018213, R01EY024698, R01EY026682, R21AG050437, New York State (N13G-275), and Foundation Fighting Blindness New York Regional Research Center Grants (PPA-1218-0751-COLU), 7T32DK00755925 and 1K01DK123199.

---

**Table S1.** Summary of BBS primary fibroblast and iPSC lines.

| **ID** | **Diagnosis** | **Affected** | **Source** | **Genotyping** | **Age** | **Sex** | **iPSC Line** |
| --- | --- | --- | --- | --- | --- | --- | --- |
| GM05948 | BBS | Yes | Coriell | BBS10<br>C91fsX95/C91fsX95 | 18 YR | Male | **BBS10A** |
| GM05950 | BBS | Yes | Coriell | BBS10<br>S303RfsX305/+ | 19 YR | Male | **BBS10B\*** |
| 1085 | BBS | Yes | Skin Biopsy | BBS1<br>M390R/M390R | 29 YR | Male | **BBS1A\*** |
| 1097 | BBS | Yes | Skin Biopsy | BBS1<br>M390R/M390R | 45 YR | Female | **BBS1B\*** |
| 1016 | Control | No | Skin Biopsy | Normal | 34 YR | Male | **Control1\*** |
| 1023 | Control | No | Skin Biopsy | Normal | 25 YR | Male | **Control2\*** |
| 1097<br>CRISPR line | Isogenic<br>control |  |  |  |  |  | **c-BBS1B** |
| 1065 | BBS | Yes | Skin Biopsy | BBS6<br>L363P/L363P | 18 YR | Male | Fibroblast<br>and iPSC |
| 1084 | BBS | Yes | Skin Biopsy | ND |  |  | Fibroblast<br>and iPSC |
| 1100 | BBS | Yes | Skin Biopsy | BBS13<br>R479H/+ | 7 YR | Female | Fibroblast |
| 1122 | BBS | Yes | Skin Biopsy | ND |  |  | Fibroblast |
| 1198 | BBS | Yes | Skin Biopsy | ND |  |  | Fibroblast |
| 1199 | BBS | Yes | Skin Biopsy | ND |  |  | Fibroblast |

\*As reported in previous published paper(35).

**Table S2.**
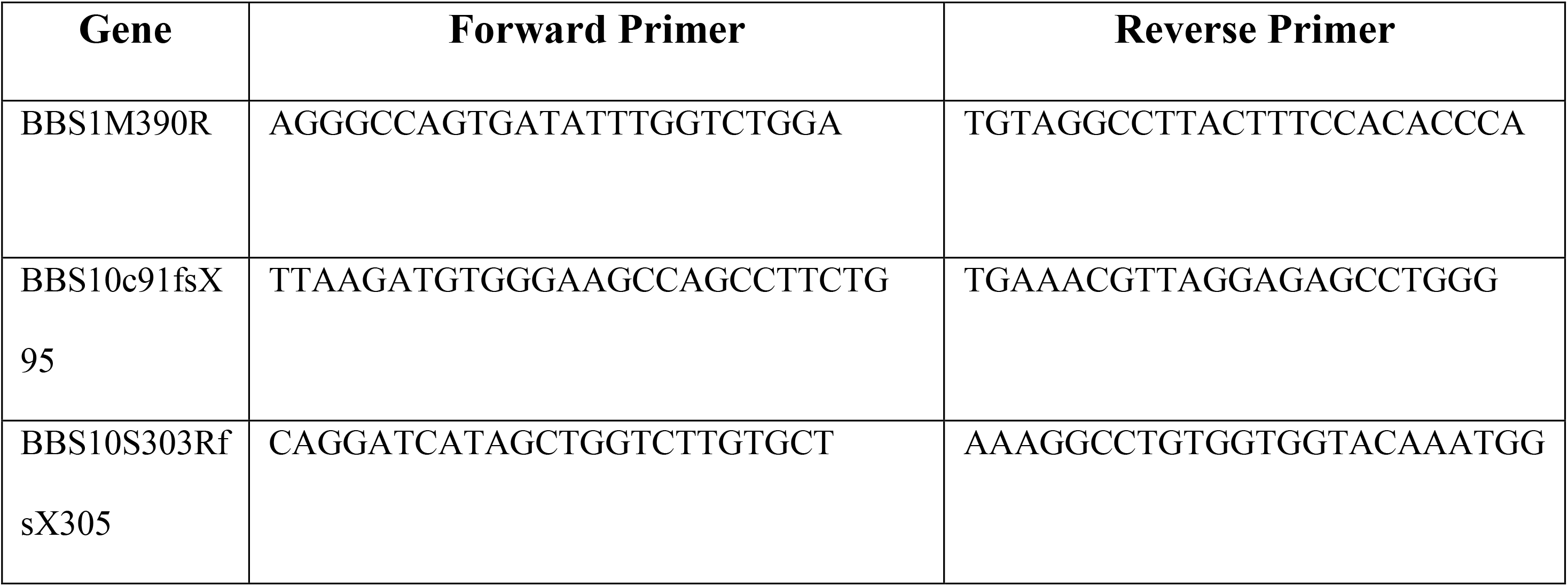
Primers for genotyping.

**Table S3.**
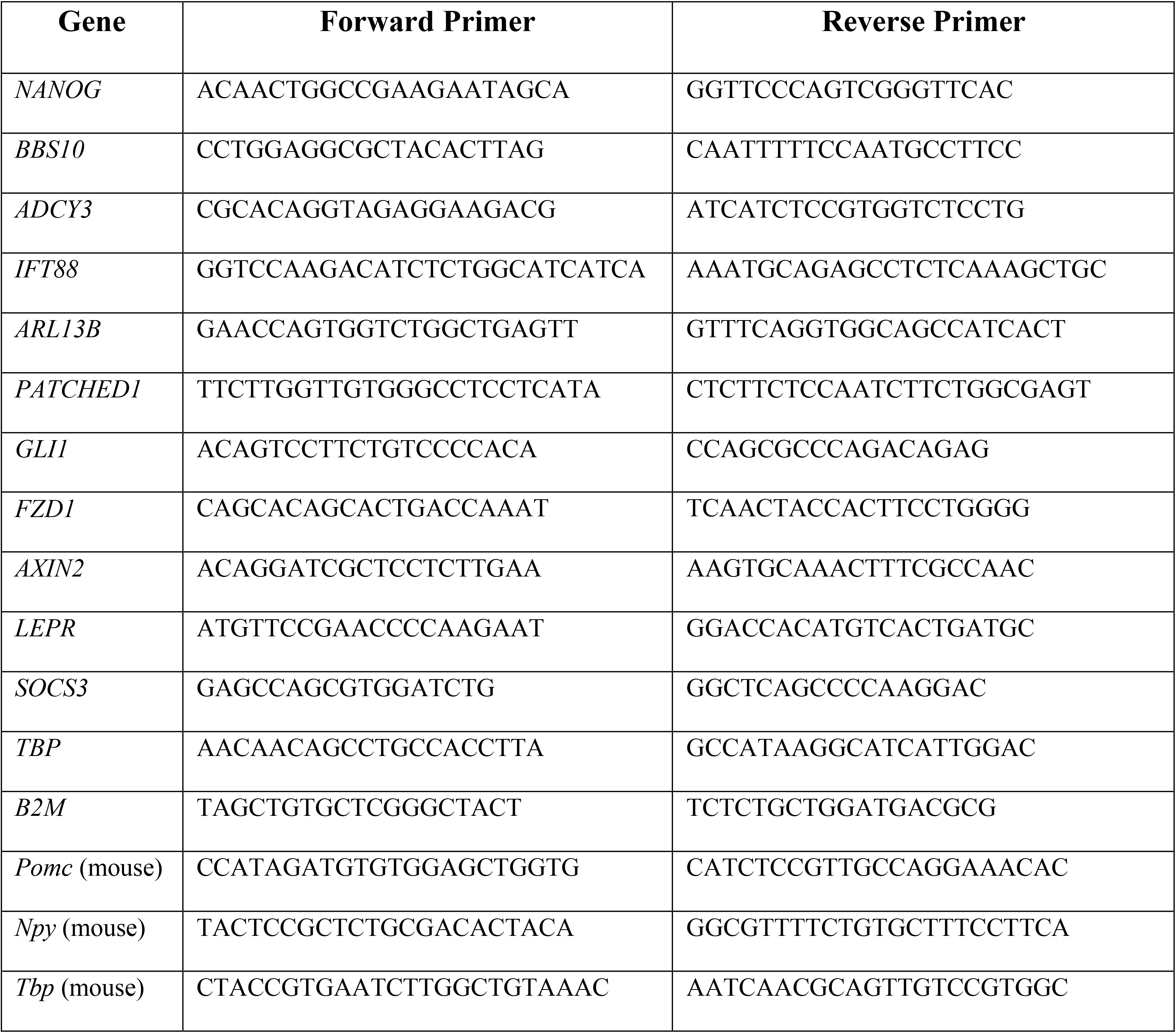
qPCR primers.

**Table S4.**
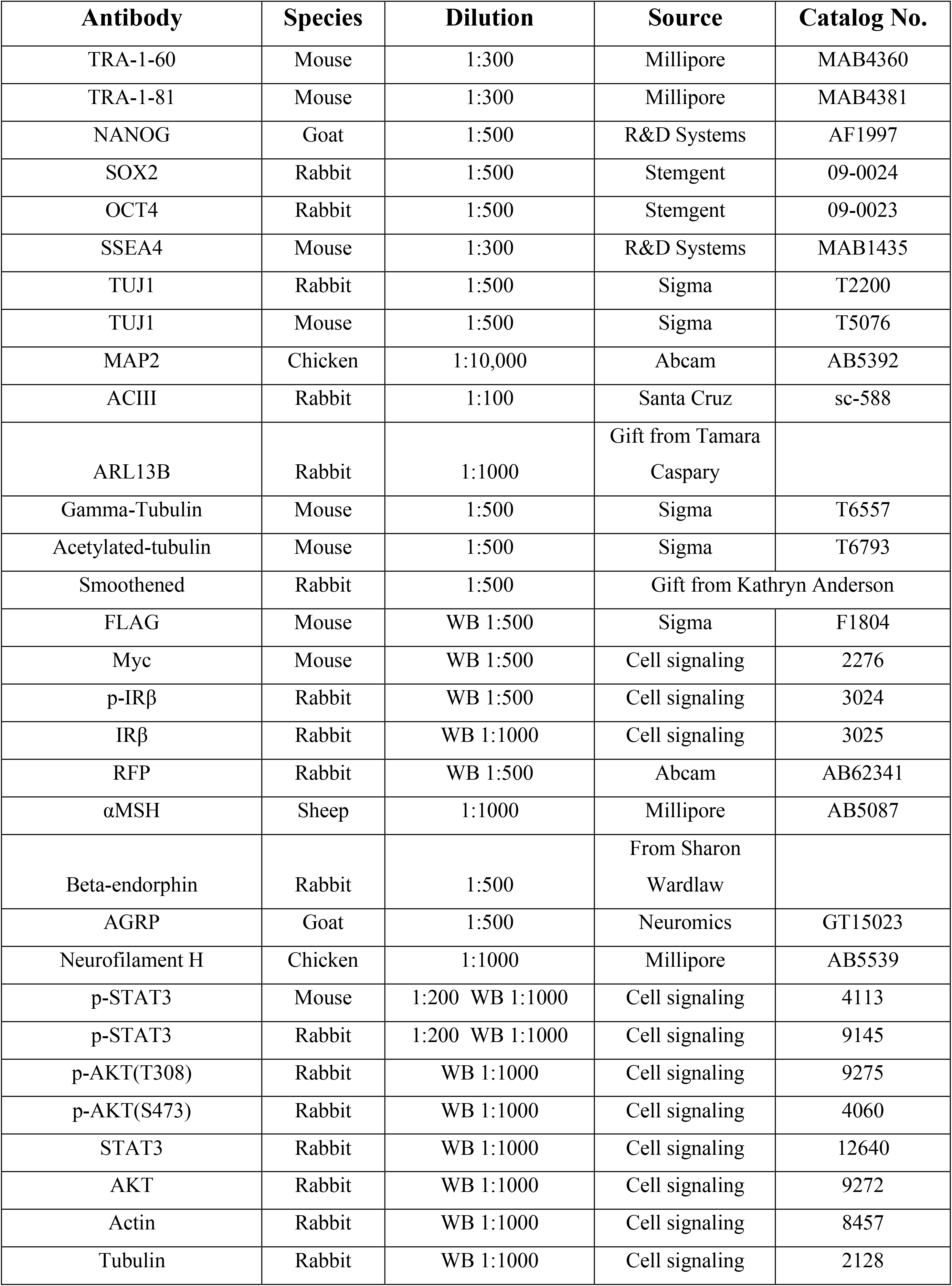
List of antibodies.

**Figure S1.**
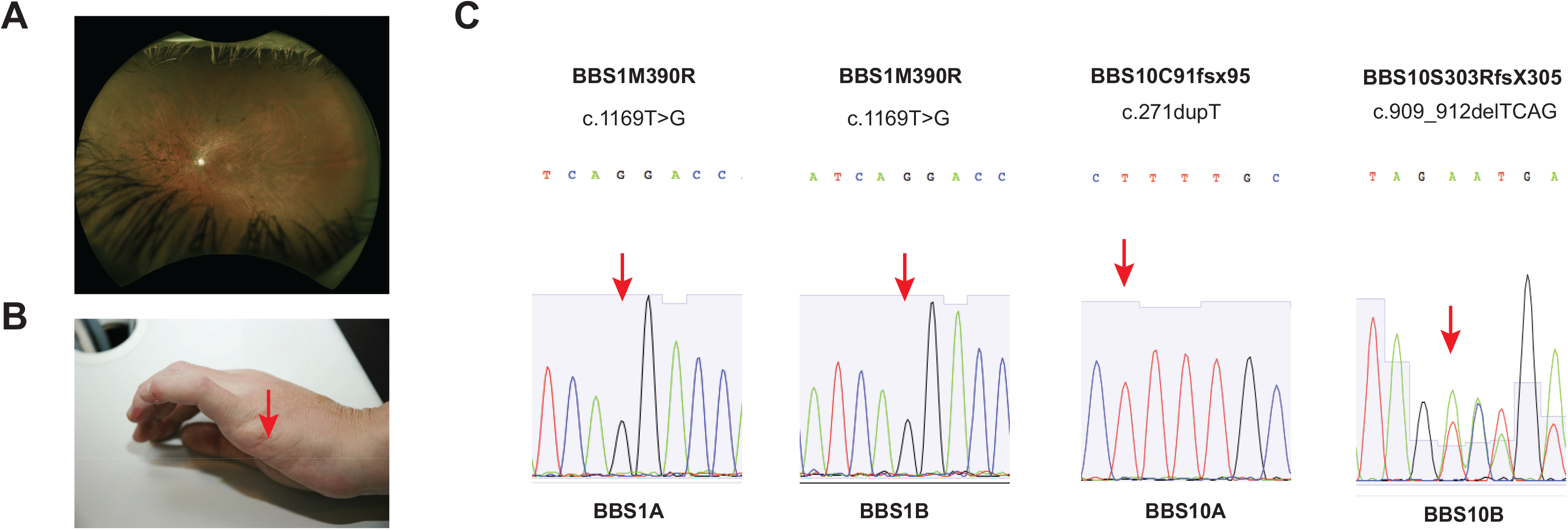
BBS patients - phenotype and genotype. (A-B) Clinical phenotypes of BBS subject *BBS1B* includes obesity, retinitis pigmentosa (A), polydactyly (B), cognitive impairment and renal cysts. Arrow points to the scar after surgery to remove the extra digit in one BBS subject. (C) Dideoxy-sequencing confirming the mutations identified by genetic testing or genome sequencing in four BBS subjects. Red arrows indicate mutation sites.

**Figure S2.**
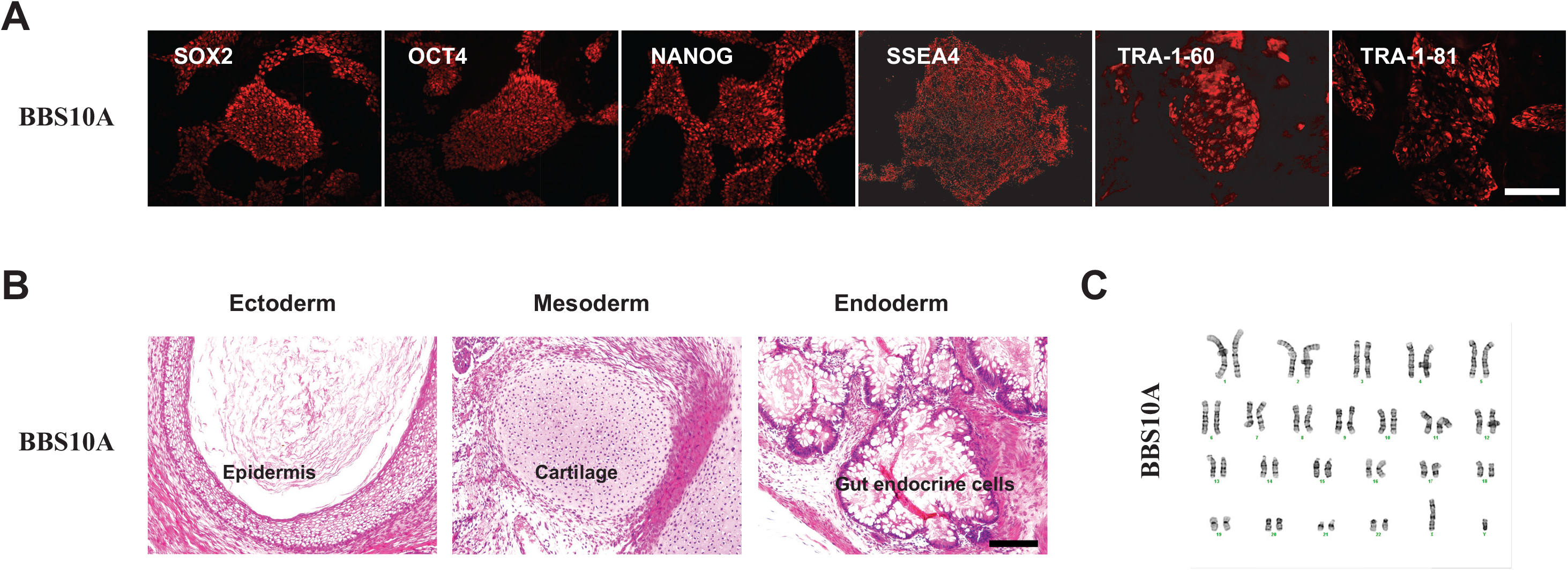
*BBS10A* iPSC line is pluripotent and has a normal karyotype. (A) Immunocytochemistry analysis of *BBS10A* iPSC line with pluripotency markers as indicated. Scale bar, 200μm; (B) H&E staining of *BBS10A* iPSC-derived teratoma sections. Cell types representing all three germ layers in iPSC-derived teratoma were identified. Scale bar, 200μm. (C) *BBS10A* iPSC had a normal karyotype.

**Figure S3.**
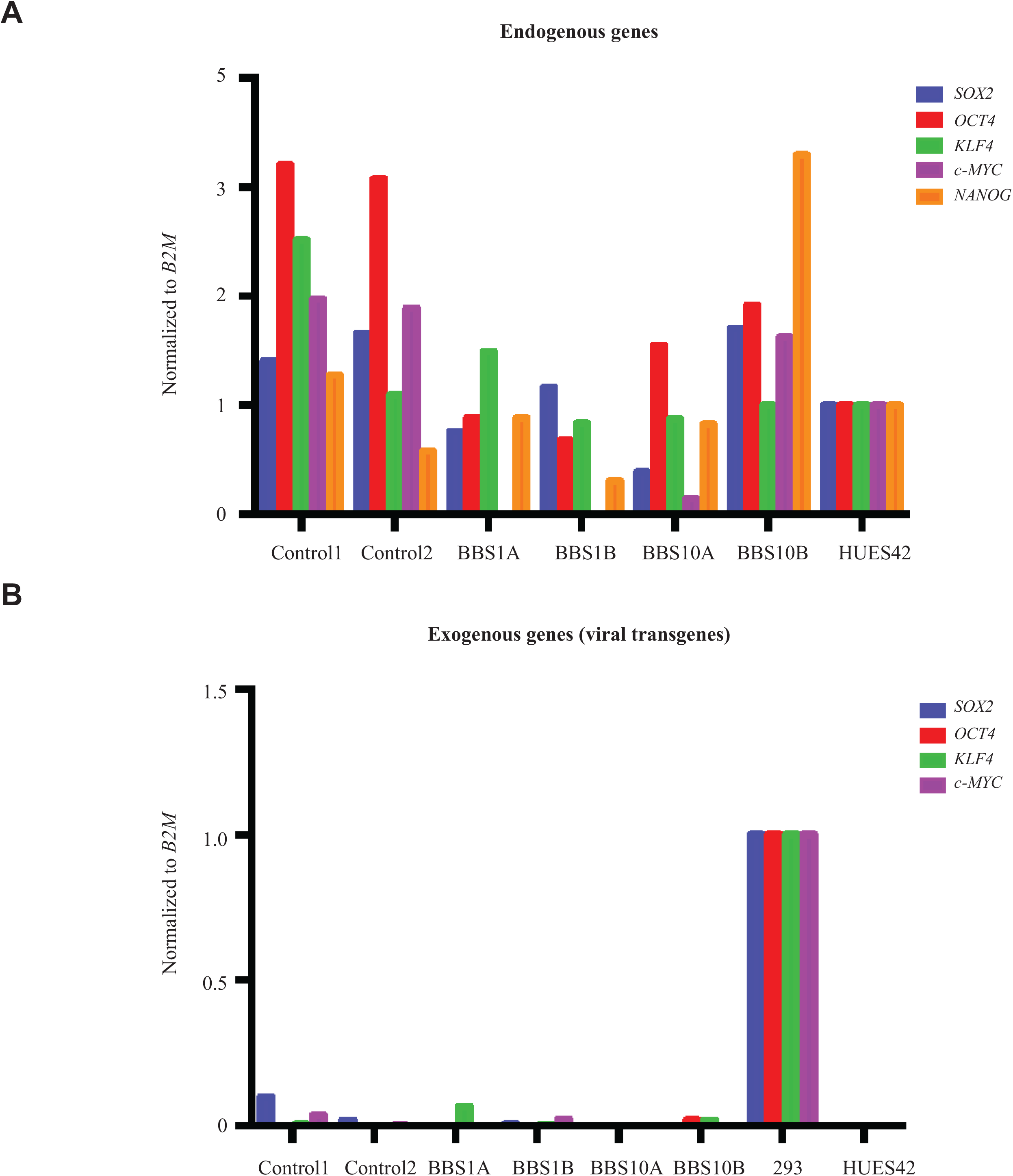
Reprogramming re-activates endogenous pluripotency genes and silences exogenous genes in iPSCs. (A) Expression of endogenous stem cell markers: *SOX2, OCT4, KLF4, c-MYC* and *NANOG* in indicated iPSC lines (passage 8-10). Human embryonic stem cell line HUES42 was used as a positive control for endogenous stem cell gene expression; (B) Expression of retroviral genes in reprogrammed iPSC as indicated. 293 cells transduced with retroviral cocktail for 48 hours were used as a positive control for viral gene expression. HUES42 was used as a negative control.

**Figure S4.**
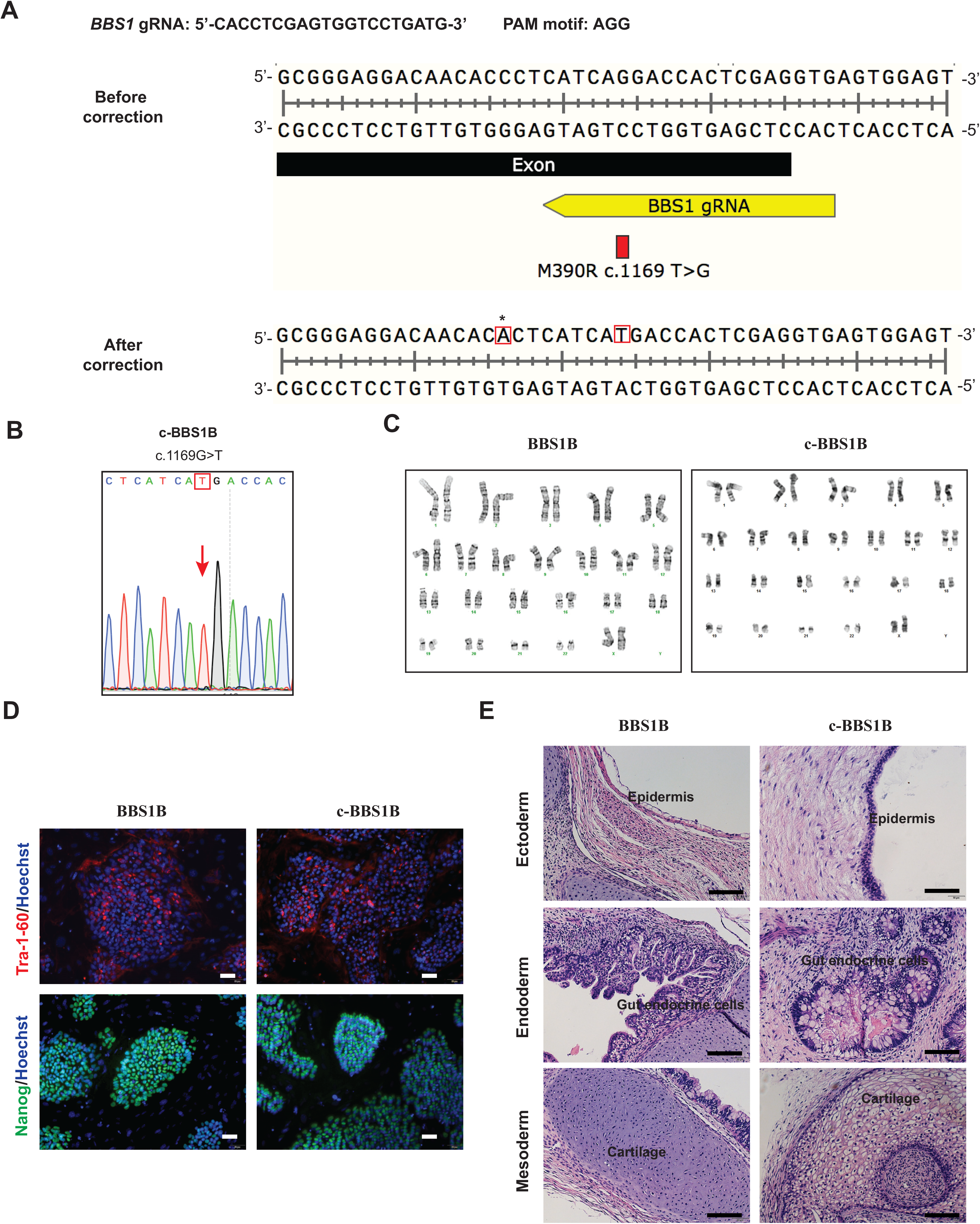
Generation of isogenic control iPSC line (*c-BBS1B*) using CRISPR/Cas9. (A) Schematic of targeting the BBS1 locus. Sequence of genomic DNA around the mutation site in *BBS1M390R* before (top) and after (bottom) CRISPR-Cas9 correction. Sequence of guide RNA (gRNA) for *BBS1 M390R* indicated in yellow. Red rectangle marks the mutation. Star marks the silent mutation introduced to avoid cutting of the repair oligo. (B) Sanger sequencing after TOPO cloning confirmed the homozygous correction of *BBS1M390R* mutation in the isogenic control line referred to as *c-BBS1B*. (C) *BBS1B* and *c-BBS1B* have normal karyotypes. (D) ICC analysis of *BBS1B* and *c-BBS1B* iPSC lines with pluripotency markers TRA-1-60 and NANOG. Scale bar, 50 μm; (E) H&E staining of *BBS1B* and *c-BBS1B* iPSC-derived teratoma sections demonstrating cell types representing all three germ layers. Scale bar, 100 μm.

**Figure S5.**
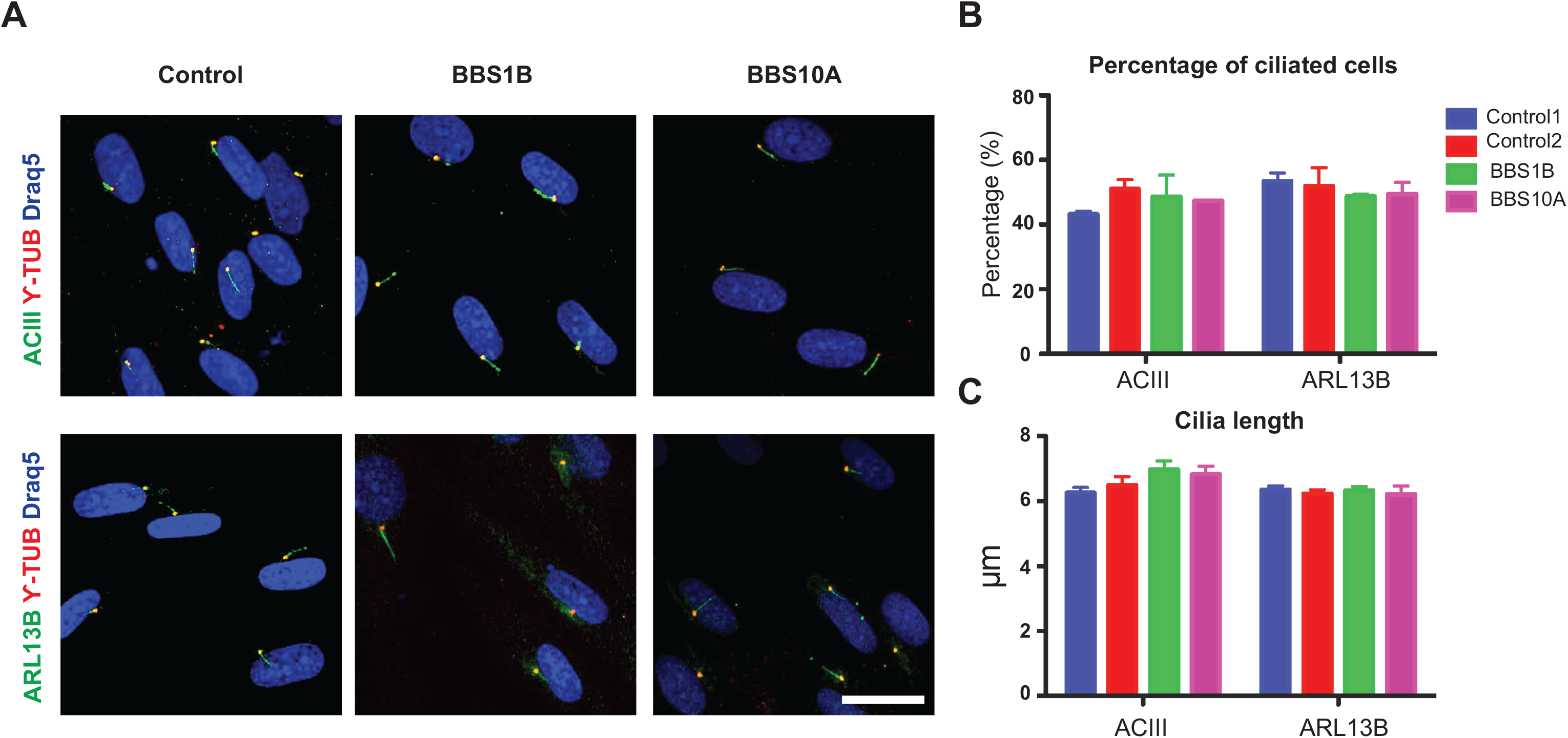
BBS mutations do not affect ciliogenesis in human fibroblasts. (A) Immunostaining of primary cilia in human fibroblasts. ACIII, ARL13B and ϒ-Tubulin are ciliary markers. Draq5 is used for nuclear staining. Scale bar, 20μm. (B-C) Quantification of percentage of ciliated cells (B) and cilia length (C) in (A).

**Figure S6.**
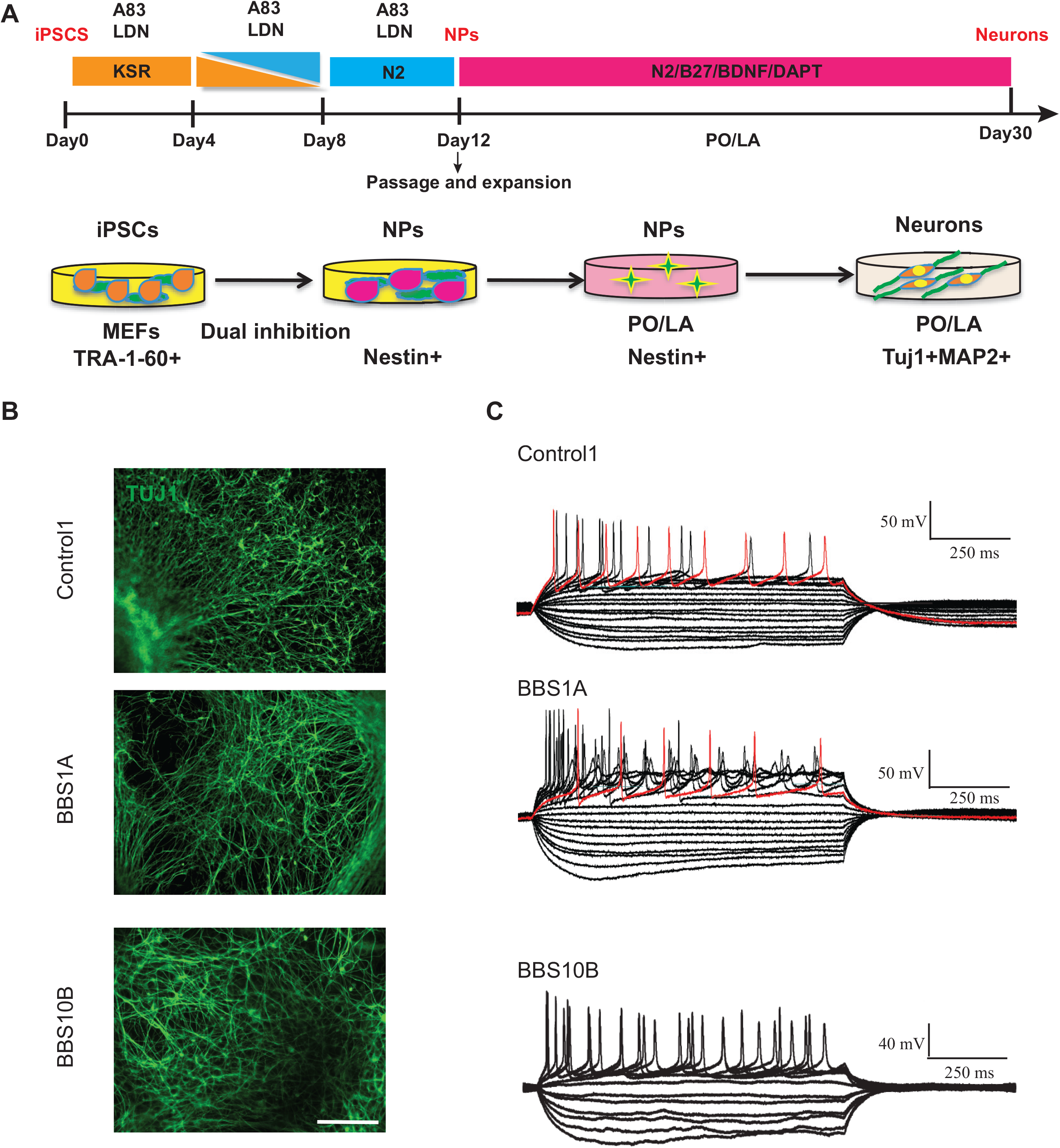
BBS mutations do not affect efficiency of neuronal differentiation or electrophysiology of iPSC-derived TUJ1+ neurons. (A) Schematic of the neuronal differentiation protocol (modified from Chambers, SM et al, 2009 (35)). TGF-β inhibitor--A83, a substitute for SB-431542, and BMP inhibitor LDN were used for dual inhibition. Day12 neuron progenitors (NPs) generated via dual SMAD inhibition were passaged onto Poly-ornithine/Laminin (PO/LA)-coated plates in N2/A83/LDN medium. Neurons were obtained around day 30 by culturing the NPs on PO/LA plates in N2/B27/BDNF/DAPT medium. (B) TUJ1 staining of Day 34 iPSC-derived neurons from control1, BBS1A and BBS10B lines. Scale bar, 200μm. (C) Whole cell patch clamp of control and BBS iPSC-derived neurons. Red trace indicates one action potential train after current stimulation.

**Figure S7.**
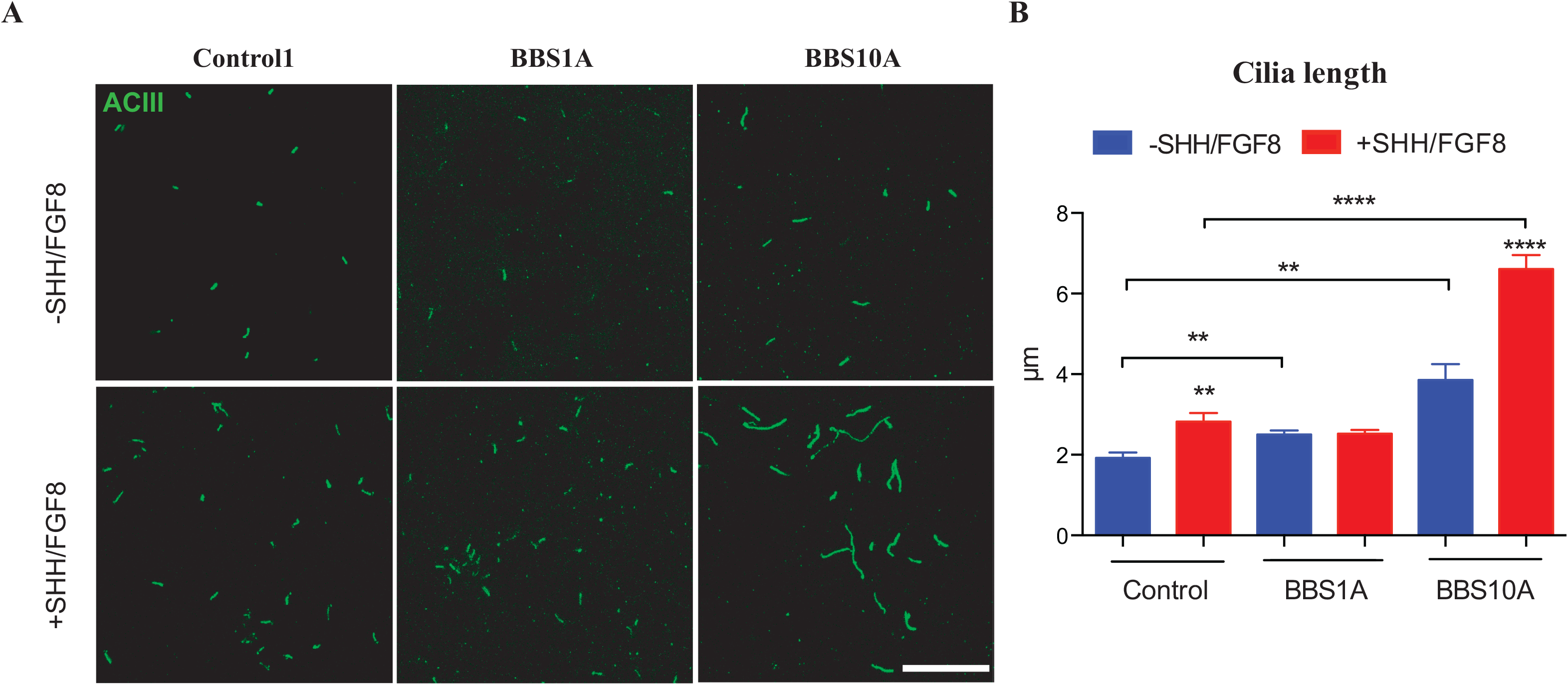
SHH/FGF8 promotes ciliogenesis in Control and *BBS10* mutant iPSC-derived neurons. (A) SHH/FGF8 treatment increased cilia length in both control and BBS iPSC-derived neurons. ACIII staining of primary cilia in Day 38 control and BBS iPSC-derived neurons. Day 35 iPSC-derived Tuj1+ neurons were treated with 0 or 100ng/ml SHH/FGF8 for 3 days. Neurons were fixed and stained with ACIII. The effect was most striking in BBS10 mutant neurons, in which “noodle-like” primary cilia were observed. Scale bar, 20μm. (B) Quantification of cilia length in (A). ** p<0.01, **** p<0.0001. Student t-test was used.

**Figure S8.**
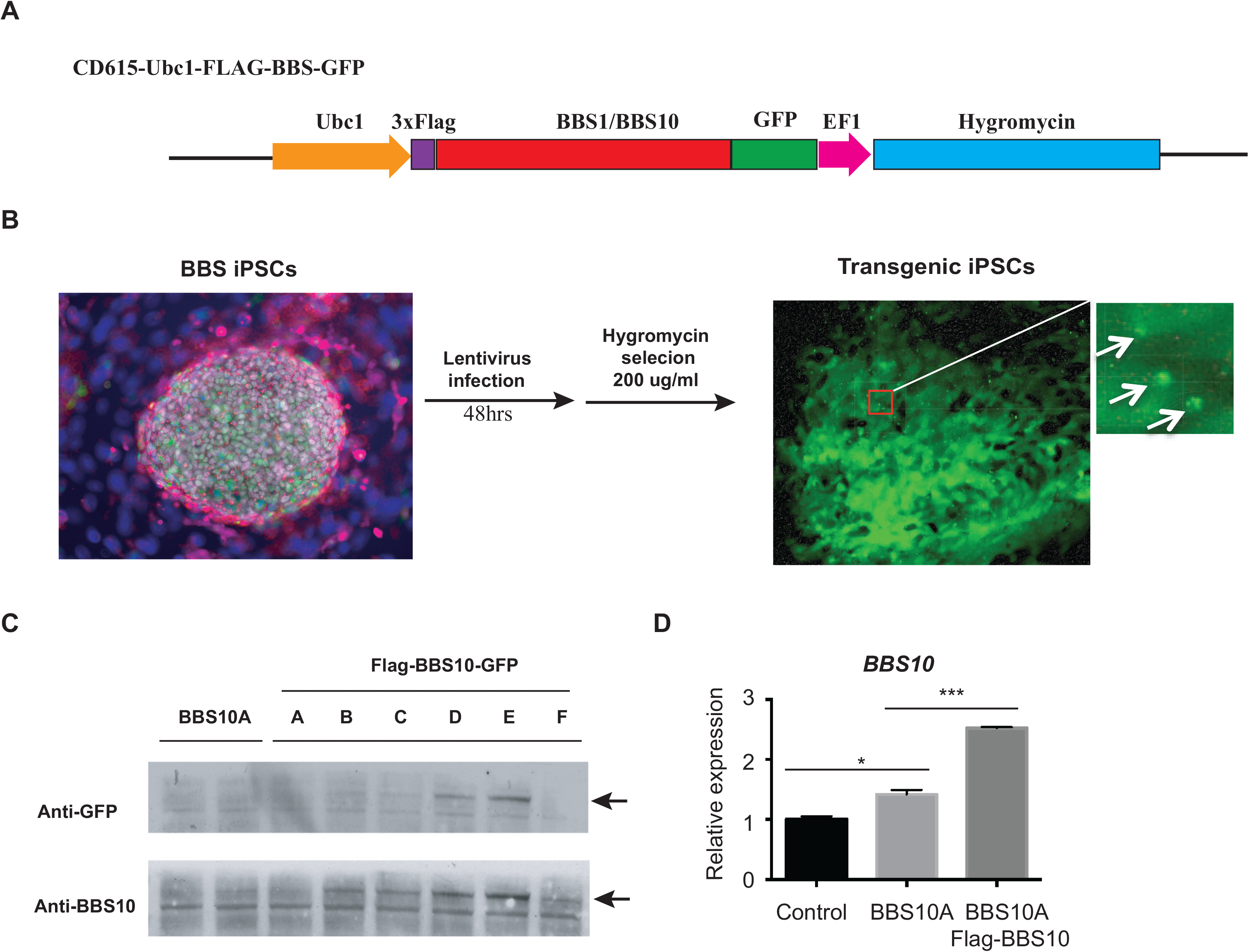
Generation of transgenic Flag-BBS1/BBS10-GFP iPSC lines. (A) Schematic of the CD615-Ubc1-FLAG-BBS-GFP lentiviral plasmid construct. (B) Experimental flows for generating BBS transgenic iPSC lines using lentivirus made from plasmid in A. BBS iPSCs were infected with lentiviral particles. 48 hrs later, the virus-containing medium was removed and hygromycin (200ug/ml) selection was applied until stable Flag-BBS10-GFP cell lines were obtained. The upper right panel shows the expression of Flag-BBS10-GFP in the transgenic *BBS10A* iPSCs, as pointed out by arrows. (C) Western blot analysis to confirm the expression of Flag-BBS10-GFP in different BBS iPSC clones. GFP and BBS10 were probed. Arrow points out Flag-BBS10-GFP band. (D) Overexpression of *BBS10* in Flag-BBS10 transgenic *BBS10A* iPSC-derived neurons. * p<0.05, ** p<0.01,*** p<0.001, student t-test.

**Figure S9.**
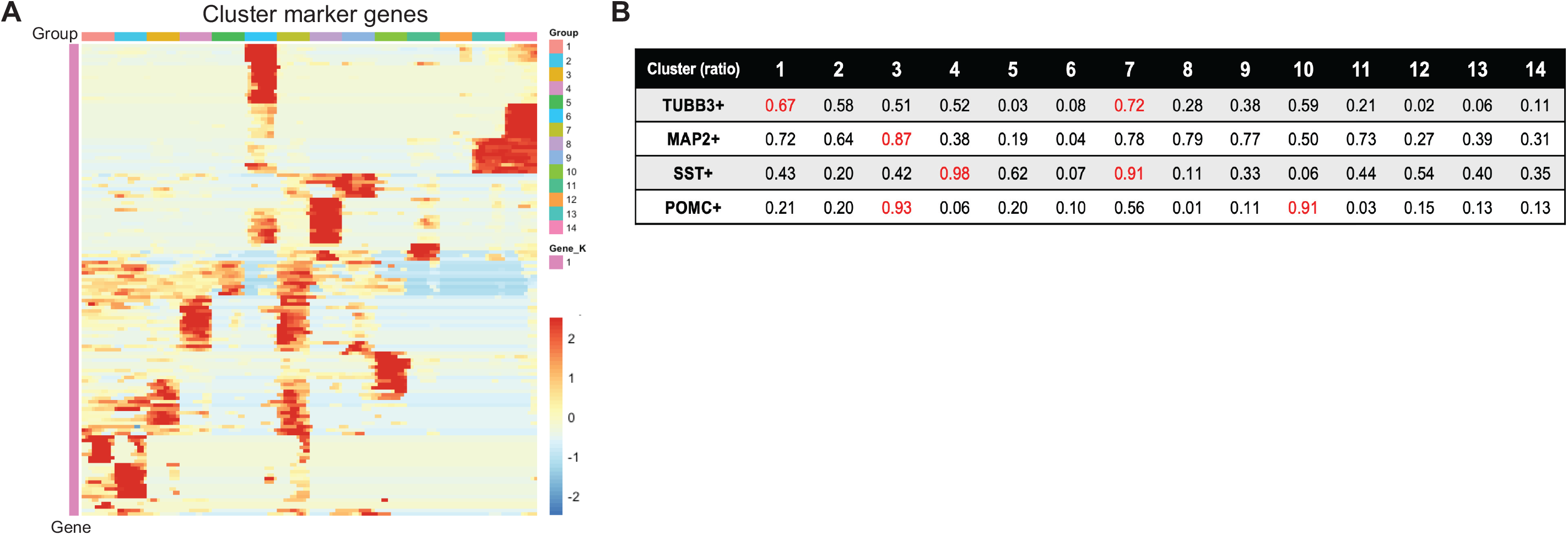
scRNA-seq of iPSC-derived hypothalamic neurons. (A) Heatmap of hierarchical clustering analysis across all 14 clusters for cluster marker gene identification. (B) Summary of ratios of TUBB3+, MAP2+, SST+ and POMC+ in all 14 clusters from *BBS1B* and *c-BBS1B* integration. Ratio per cluster = positive cells/total cells. Cut off: median expression level.

**Figure S10.**
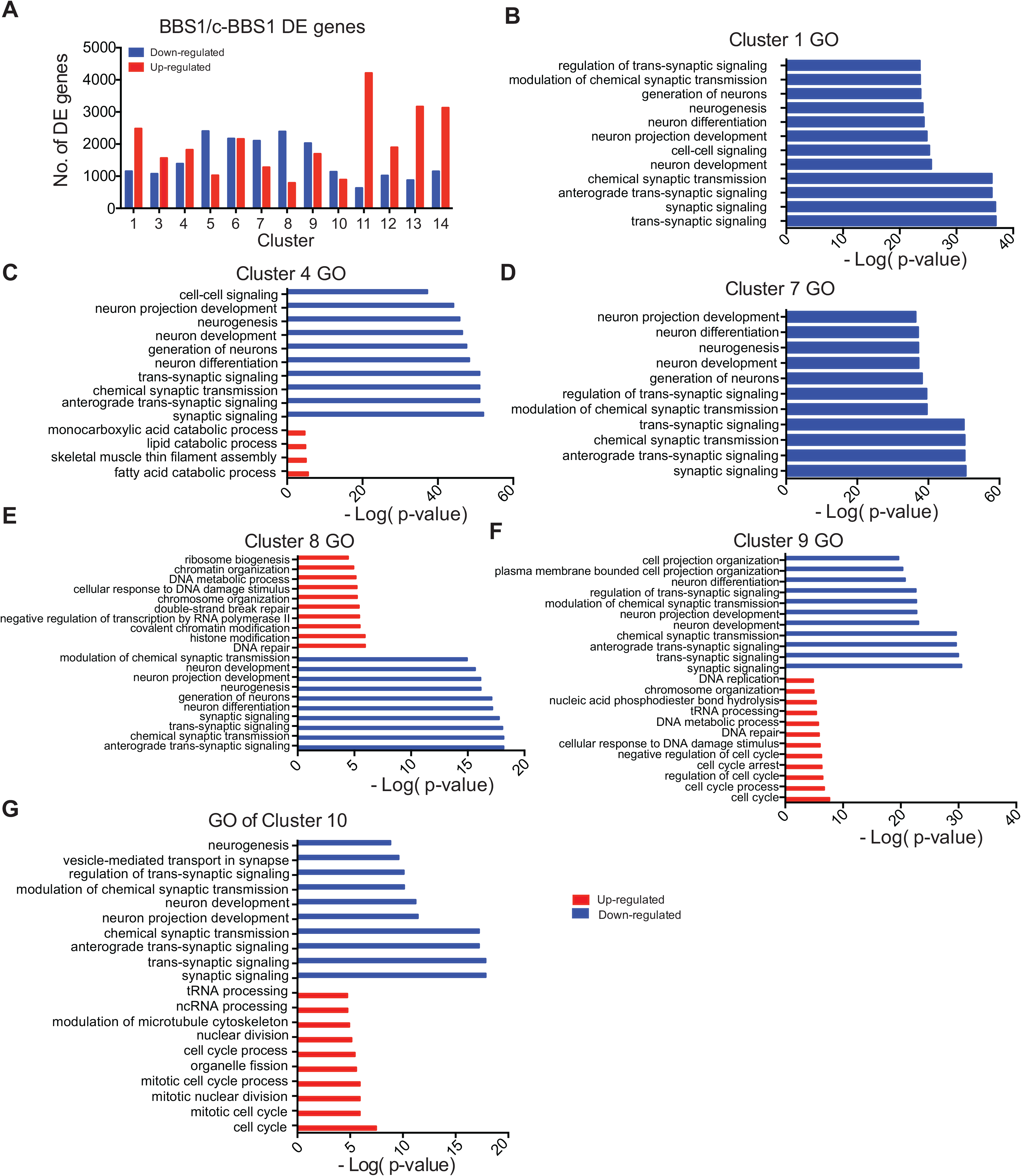
Gene ontology analysis of differentially expressed genes in neuronal clusters. (A) Distribution of differentially expressed genes in all 14 clusters. No. of up- and down-regulated expressed genes (*BBS1B* vs *c-BBS1B*, P<0.05) were recorded. (B-G) Gene ontology for biological process analysis for clusters 1, 4, 7-10. Adjusted p-value was used for this plot.

**Figure S11.**
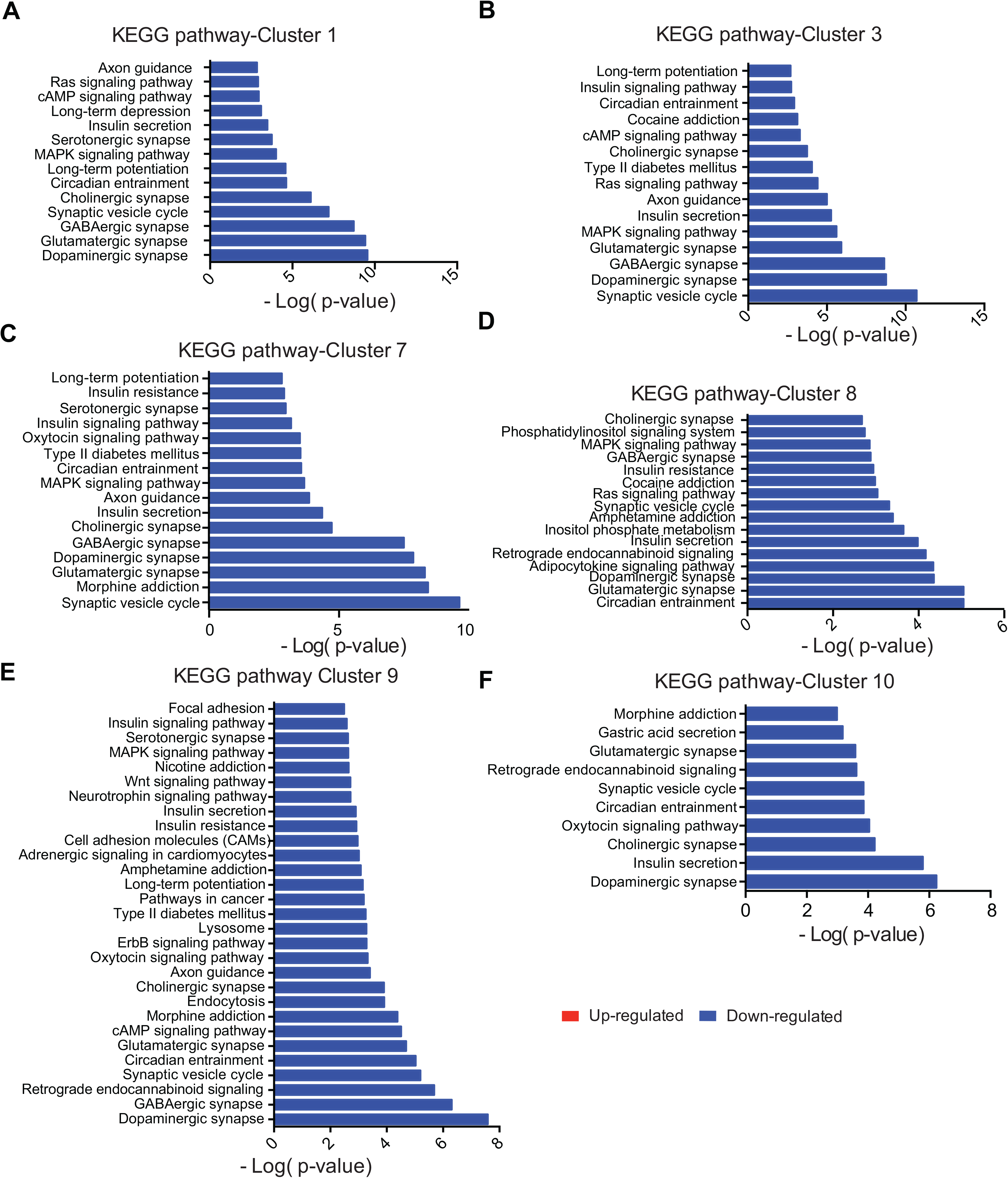
KEGG pathway analysis of differentially expressed genes in neuronal clusters. KEGG pathway analysis of both up- and down-regulated differentially expressed genes (*BBS1B* vs *c-BBS1B*) for clusters 1, 3, 7-10. Adjusted p-values were used for this plot.

**Figure S12.**
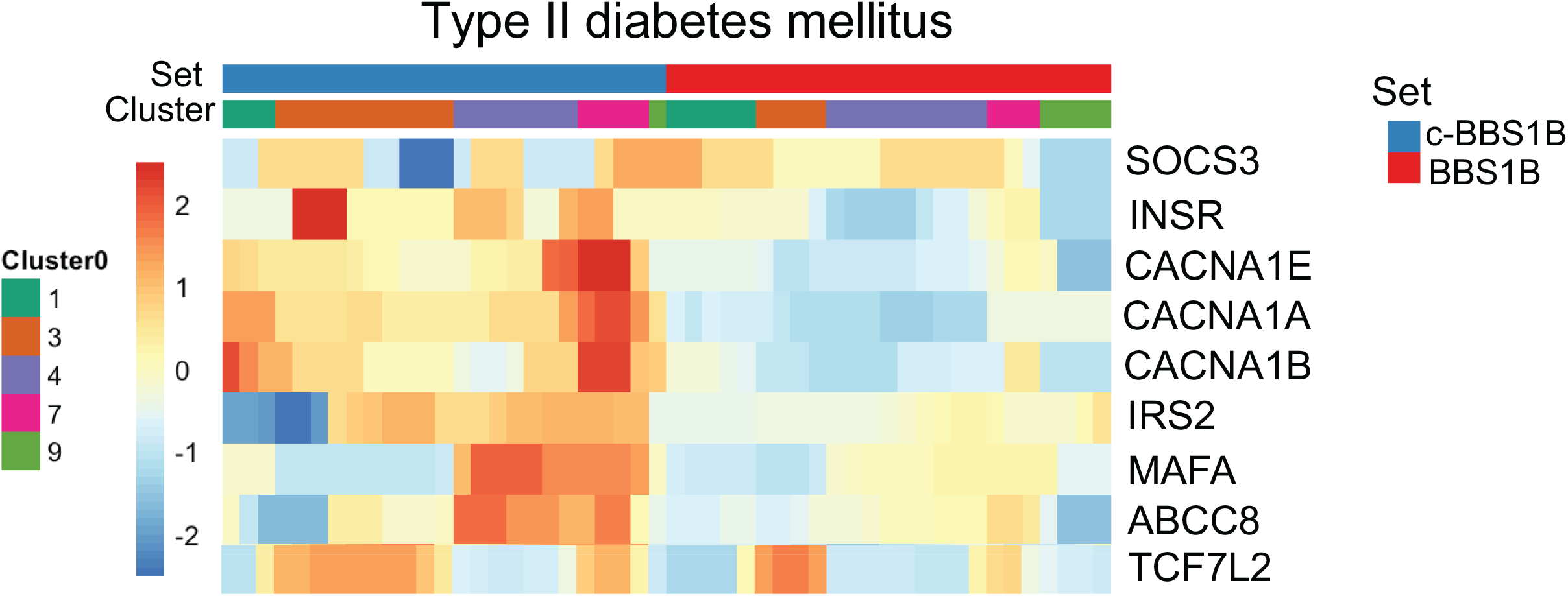
scRNA-seq reveals that *BBS1M390R* mutation decreases expression of genes related to type II diabetes mellitus pathway. Heatmap of genes involved in type II diabetes mellitus from KEGG pathway analysis. Clusters1, 3, 4, 7 and 9 of *BBS1B* and *c-BBS1B* iPSC-derived hypothalamic neurons (p<0.05) were included.

**Figure S13.**
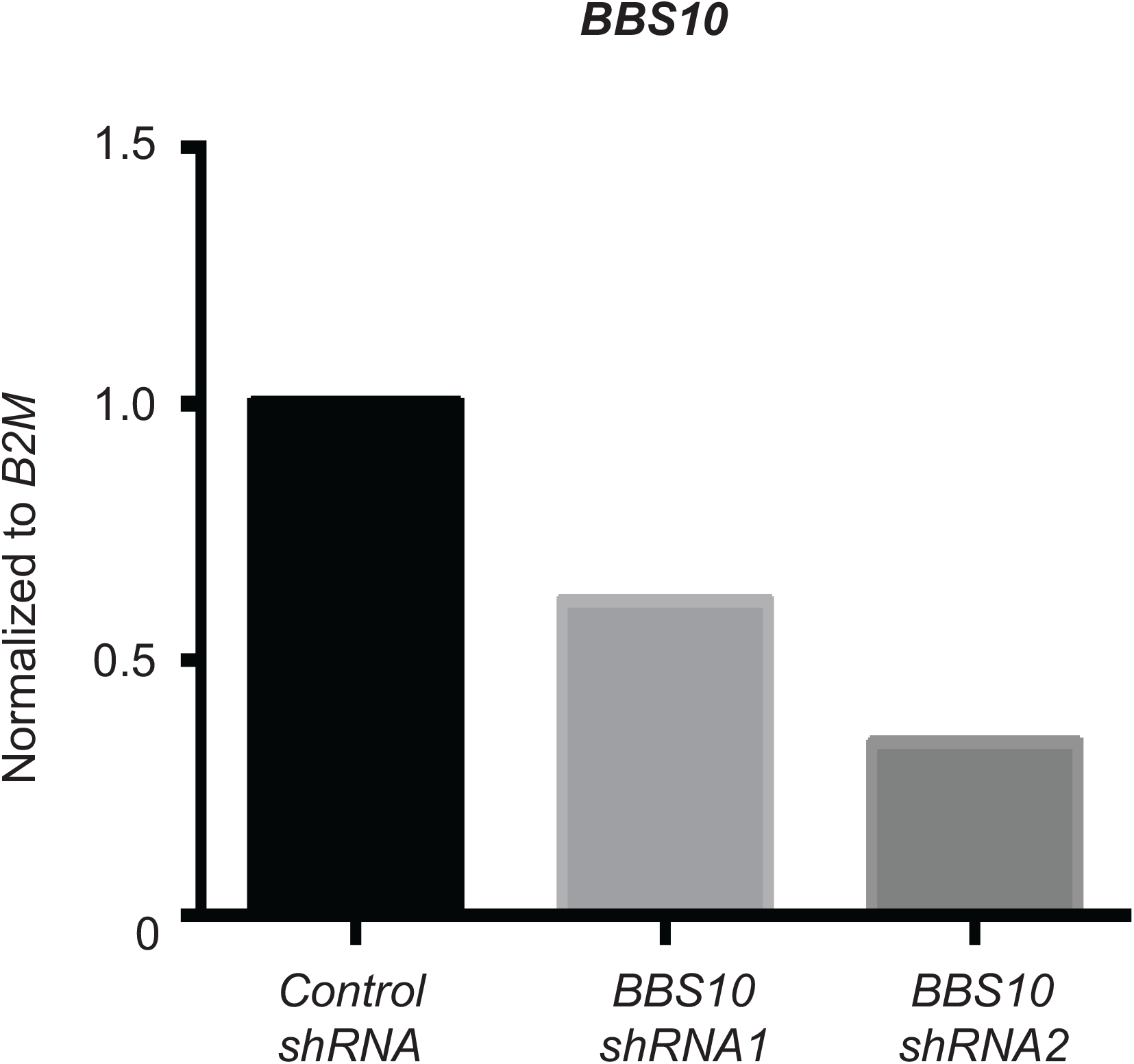
Knockdown of *BBS10* in control iPSCs using lentiviral shRNA. Control iPSCs were infected with control and two *BBS10* lentiviral shRNAs for 48hrs. These iPSC were further treated with 1μg/ml puromycin selection until puromycin resistant iPSC were obtained. Expression of *BBS10* in shRNA knockdown iPSC lines were determined by RT-qPCR. N=1.

**Figure S14.**
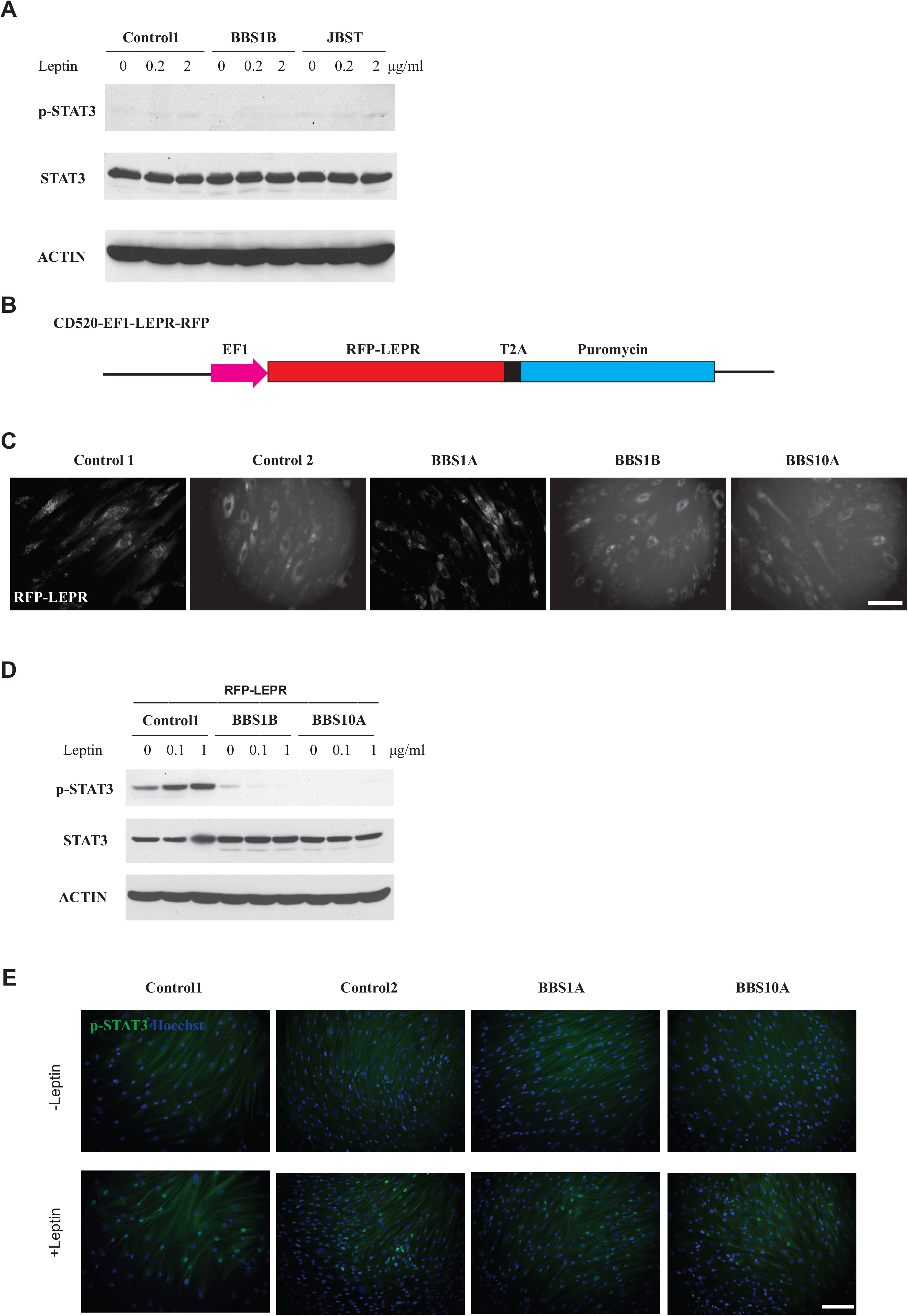
BBS mutations impair leptin signaling in RFP-LEPR transgenic human fibroblasts. (A) Western blot analysis of leptin signaling in human control, BBS and JBST fibroblasts. Human fibroblasts were serum starved overnight and exposed to 0, 0.2 and 2 μg/ml leptin for 30min. (B) Diagram of the lentiviral RFP-LEPR construct.(C) Live cell imaging shows RFP expression in RFP-LEPR transgenic control and BBS fibroblast lines as indicated. Scale bar: 100 μm; (D) Western blot analysis of leptin signaling in RFP-LEPR transgenic human fibroblasts. *Control*, *BBS1B* and *BBS10A* RFP-LEPR transgenic fibroblasts were included. Cells were exposed to 0, 0.1 and 1 μg/ml leptin for 30min. p-STAT3, STAT3 and ACTIN were analyzed from cell lysates. (E) Staining of p-STAT3 in control and BBS RFP-LEPR transgenic human fibroblasts. Human fibroblasts were fasted overnight and exposed to 1 μg/ml leptin for 30min. Cells were fixed and stained with p-STAT3 and Hoechst. Scale bar, 200μm.

**Figure S15.**
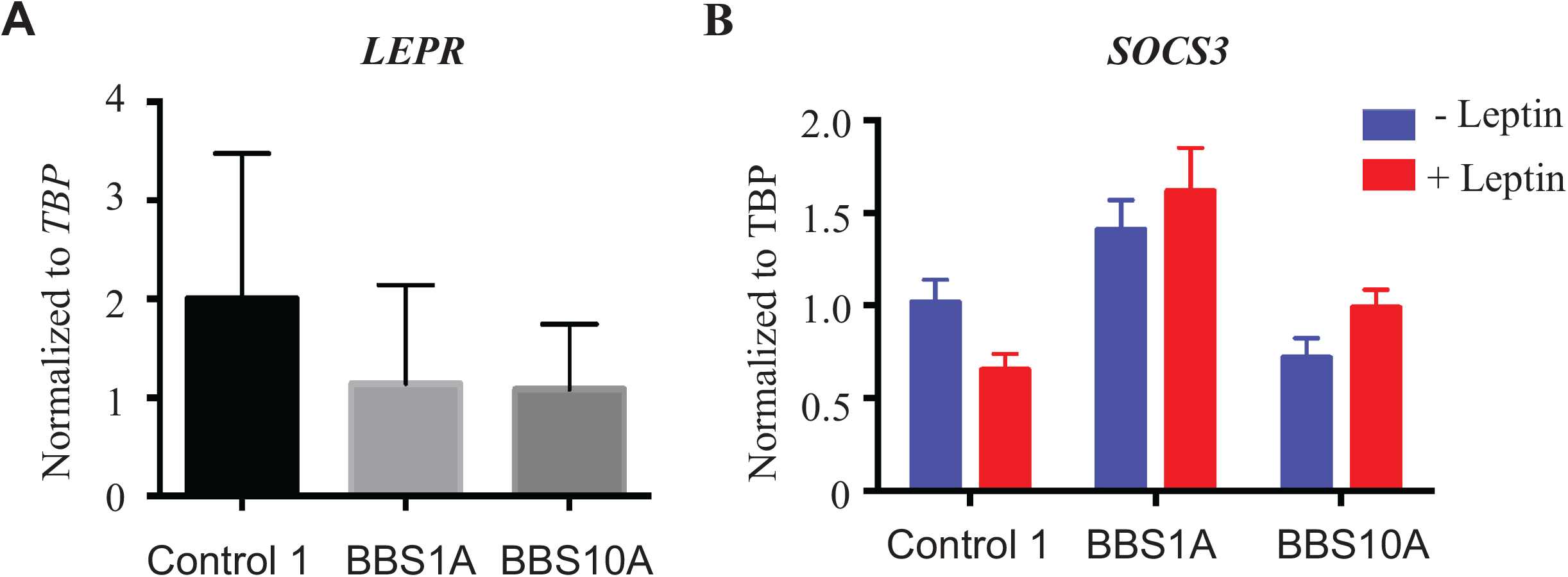
Expression of leptin signaling molecules in iPSC-derived neurons. (A) Expression of *LEPR* in iPSC-derived neurons; (B) Expression of *SOCS3* in *Control* and *BBS* iPSC-derived neurons in the presence or absence of leptin (1ug/ml) for 24hrs.

